# Shared retinoic acid responsive enhancers coordinately regulate nascent transcription of *Hoxb* coding and non-coding RNAs in the developing mouse neural tube

**DOI:** 10.1101/2022.08.30.505933

**Authors:** Zainab Afzal, Jeffrey Lange, Christof Nolte, Sean McKinney, Christopher Wood, Ariel Paulson, Bony De Kumar, Jay Unruh, Brian D. Slaughter, Robb Krumlauf

## Abstract

Signaling pathways regulate the patterns of *Hox* gene expression that underlie their functions in specification of axial identity. Little is known about the properties of *cis*-regulatory elements and underlying transcriptional mechanisms that integrate graded signaling inputs to coordinately control *Hox* expression. Here we optimized single molecule fluorescent *in situ* hybridization (smFISH) technique with probes spanning introns to evaluate how three shared retinoic acid response element (RARE)-dependent enhancers in the *Hoxb* cluster regulate patterns of nascent transcription *in vivo* at the level of single cells in wild type and mutant embryos. We predominately detect nascent transcription of only a single *Hoxb* gene in each cell, with no evidence for simultaneous co-transcriptional coupling of all or specific subsets of genes. Single and/or compound RARE mutations indicate each enhancer differentially impacts global and local patterns of nascent transcription, suggesting that selectivity and competitive interactions between these enhancers is important to robustly maintain the proper levels and patterns of nascent *Hoxb* transcription. This implies rapid and dynamic regulatory interactions potentiate transcription of genes through combined inputs from these enhancers in coordinating the RA response.

## Introduction

*Hox* genes encode a conserved family of transcription factors (TFs) that play an important role in regulating regional identity and diversification of tissues along the anterior-posterior (A-P) axis during development and in the evolution of a wide variety of animal species, from invertebrates to vertebrates (Lewis, 1978; Duboule and Dolle, 1989; Graham et al., 1989; McGinnis and Krumlauf, 1992; Carroll, 1995; Mallo et al., 2010; Martin et al., 2016; Arendt, 2018; He et al., 2018). *Hox* genes are arranged in tightly linked clusters and conserved features in the organization of these chromosomal clusters results in the generation of highly ordered patterns of *Hox* expression and function along the A-P axis of developing embryos (Lewis, 1978; Duboule and Dolle, 1989; Graham et al., 1989; Serano et al., 2016). Through coordinate regulation of their spatial and temporal patterns of expression, *Hox* genes lay down a combinatorial code that is a key component of the regulatory hierarchy which specifies the regional properties of tissues and modulates morphological diversity. Disruptions in the expression domains of *Hox* genes result in homeotic transformations and altered morphogenesis (Balkaschina, 1929; Bridges and Dobzhan, 1933; Garber et al., 1983; Hafen et al., 1984; Pultz et al., 1988; Merrill et al., 1989; Maconochie et al., 1996; Philippidou and Dasen, 2013; Quinonez and Innis, 2014). Hence, during development the levels and patterns of *Hox* gene expression along the A-P axis must be precisely regulated in space and time for proper elaboration of the basic body plan. In early embryogenesis, graded cues from major signaling pathways, such as retinoic acid (RA), Wnt (Wingless related integration site) and Fgf (Fibroblast growth factors), play a key role in organizing the establishment and maintenance of patterns of *Hox* gene expression that underlie their functions in specification of axial identity and assignment of cell fate (Simeone et al., 1990; Pownall et al., 1996; Bel-Vialar et al., 2002; Diez del Corral and Storey, 2004; Deschamps and van Nes, 2005; Schilling et al., 2012; Neijts et al., 2016; Deschamps and Duboule, 2017; Darras et al., 2018; Frank and Sela-Donenfeld, 2019; Nolte et al., 2019). Therefore, it is important to understand the properties of *cis*-regulatory elements and transcriptional mechanisms that interpret and integrate these graded signaling inputs to coordinately control the dynamic expression of *Hox* genes. These features make the clustered *Hox* genes a good paradigm for investigating transcriptional processes that underlie coordinate and combinatorial regulation of genes *in vivo* during development.

Regulatory analyses in invertebrate and vertebrate animal model systems have revealed that a wide variety of mechanisms provide inputs that serve to coordinate patterns of *Hox* expression at the level of transcription (de Laat and Duboule, 2013; Gregor et al., 2014; Gentile and Kmita, 2018; Afzal and Krumlauf, 2022; Batut et al., 2022; Gaskill and Harrison, 2022). Regulation of clustered *Hox* genes in vertebrate systems has been linked to diverse types of *cis*-regulatory elements (CREs) and differences in chromatin states, epigenetic modifications, and chromosome topology/3D genome conformations (Sharpe et al., 1998; Chambeyron and Bickmore, 2004; Tarchini and Duboule, 2006; Gonzalez et al., 2007; Soshnikova and Duboule, 2009; Noordermeer et al., 2011; Mazzoni et al., 2013; Ahn et al., 2014; Noordermeer et al., 2014; De Kumar et al., 2015; Narendra et al., 2015; Narendra et al., 2016; Neijts et al., 2016; Parker and Krumlauf, 2017; Qian et al., 2018; Rodríguez-Carballo et al., 2019). Among the CREs, enhancers play an important role in modulating the activation and/or maintenance of transcription of *Hox* genes through their ability to interpret graded cues from signaling pathways and to integrate dynamic combinations of TFs to control gene expression patterns in a spatio-temporal and tissue-specific manner (Marshall et al., 1994; Studer et al., 1994; Tümpel et al., 2002; Berlivet et al., 2013; Delpretti et al., 2013; Paris et al., 2013; Crocker et al., 2015; Heinz et al., 2015; Parker and Krumlauf, 2017; Henriques et al., 2018; Nolte et al., 2019; Parker et al., 2019; Choi et al., 2021; Kreibich et al., 2022). Studies in mice, in a variety of different tissues, have demonstrated that there are multiple enhancers embedded within and flanking the *Hox* clusters which can exhibit overlapping activities (shadow enhancers), selective and competitive preferences for target genes, and they can regulate both near adjacent genes or act more globally on multiple genes in a cluster (shared enhancers) (Oosterveen et al., 2003a; Scotti and Kmita, 2012; Tschopp et al., 2012; Andrey et al., 2013; Berlivet et al., 2013; Nolte et al., 2013; Ahn et al., 2014; Qian et al., 2018). The diverse roles of enhancers in coordinate regulation of *Hox* genes, in conjunction with the compact nature and relatively high density of coding and non-coding transcriptional units in a *Hox* cluster, raises fundamental questions about the specificity of enhancers and how they locate and distinguish between their target loci (Sanyal et al., 2012; Long et al., 2016; Furlong and Levine, 2018; Schoenfelder and Fraser, 2019; Zeitlinger, 2020; Afzal and Krumlauf, 2022; Batut et al., 2022; Gaskill and Harrison, 2022; Levo et al., 2022).

Models such as linking (Morcillo et al., 1997), tracking (Kong et al., 1997), and looping (Dunn et al., 1984; Deng et al., 2012) have been proposed to explain how enhancers can locate and activate gene promoters. It is unclear whether there are optimal enhancer-promoter distances and if such preferences are linked to regulation of spatio-temporal or tissue specific activities in development, as experimental changes in enhancer-promoter proximity have resulted in conflicting observations (Bartman et al., 2016; Benabdallah et al., 2019). Recent studies have provided evidence for rapid dynamics in interactions between enhancers and their target promoters over a wide range of distances, which has led to a revision of older models postulating stable long-term enhancer-promoter interactions in favor of models based on dynamic looping (Bothma et al., 2014; Gregor et al., 2014; Furlong and Levine, 2018; Liu and Tjian, 2018; Mir et al., 2018; Zhang and Tjian, 2018; Berrocal et al., 2020; Eck et al., 2020; Zuin et al., 2022). Furthermore, there is evidence for the occurrence of dynamic co-transcriptional hubs containing shared pools of components of the general transcription machinery and upstream activators (Tsai et al., 2017; Furlong and Levine, 2018; Mir et al., 2018; Tsai et al., 2019). In *Drosophila*, many shared enhancers of developmental genes have recently been shown to co-transcriptionally couple activity of their target genes in living embryos, even if those genes are separated by large distances, by integrating independent inputs from tethering elements and topologically associated domains (TADs) (Levo et al., 2022). This mode of genome organization creates what are referred to as “topological operons” for coordinate and co-dependent transcriptional regulation of multiple genes by shared enhancers. However, whether shared enhancers in mammals and other vertebrates also tend to regulate their target genes by similar co-transcriptional coupling mechanisms or depend upon stochastic and independent regulation of each target gene, is unknown.

The mouse *Hoxb* cluster provides a good context for investigating the *in vivo* properties and roles of shared enhancers in coordinate regulation of *Hox* gene expression in response to RA signaling. The *Hoxb1-b9* coding genes and several non-coding RNAs span a ∼100 kb region of chromosome 11 (Fig. 1A), while *Hoxb13* is located ∼70 kb downstream of *Hoxb9,* separated by a region of non-coding sequences. These genes are located within a single TAD, encompassing many sub-TADs (Dixon et al., 2012; Ke et al., 2017). Dynamic RA gradients play a central role in directly activating *Hoxb* genes in the developing hindbrain and spinal cord (Bel-Vialar et al., 2002; Oosterveen et al., 2003a; Sirbu et al., 2005; Nolte et al., 2019; Krumlauf and Wilkinson, 2021) and RA signaling coordinately modulates their expression in many other tissues, such as hematopoietic stem cells (HSCs), axial elongation and endoderm (Deschamps and van Nes, 2005; Niederreither and Dolle, 2008; Bertrand et al., 2011; Rhinn and Dolle, 2012; Zaffran and Kelly, 2012; Nolte et al., 2013; Deschamps and Duboule, 2017; Qian et al., 2018). The direct transcriptional responses of the *Hoxb* cluster to graded RA signaling are integrated in part through a series of retinoic acid response elements (RAREs) embedded within and flanking the cluster (Marshall et al., 1994; Studer et al., 1994; Dupe et al., 1997; Gould et al., 1998; Houle et al., 2003; Nolte et al., 2003; Oosterveen et al., 2003b; Nolte et al., 2013; Ahn et al., 2014; Nolte et al., 2019). For example, three conserved RAREs (*DE*, *B4U*, and *ENE*) are components of RA-dependent enhancers present in a 15 kb region in the middle of the cluster, which also contains transcription units for *Hoxb4, Hoxb5, mir10a* and two long non-coding (lnc) RNAs implicated in regulation of *Hoxb* genes (*Hobbit* and *HoxBlinc*) (De Kumar et al., 2015; De Kumar and Krumlauf, 2016; Deng et al., 2016; Degani et al., 2021) (Fig. 1A). These three shared enhancers potentiate global *Hoxb* responses to RA by regulating multiple coding and non-coding transcripts (Gould et al., 1997; Gould et al., 1998; Sharpe et al., 1998; Oosterveen et al., 2003a; Oosterveen et al., 2003b; Ahn et al., 2014). The *DE-RARE* is an essential element in an RA-dependent enhancer, which is subject to epigenetic modifications, and required for coordinating global regulation of multiple *Hoxb* coding and *lncRNAs* in HSCs and the developing neural tube, as indicated by arrows in Fig. 1A (Ahn et al., 2014; De Kumar et al., 2015; Qian et al., 2018). The *ENE-RARE* regulates *Hoxb4* and *Hoxb3* and transgenic analyses suggest some degree of functional overlap with the *DE-RARE* in regulating the RA response of other genes in the cluster (Gould et al., 1997; Ahn et al., 2014). This high density of enhancers and transcription units, in such a small region (Fig. 1A), raises questions regarding how these enhancers participate in short and long-range regulation of multiple genes, how and which targets are selected, the dynamics of the process and whether there is co-transcriptional coupling of regulated transcriptional units. In addition, whether these three RARE-dependent enhancers exert their effects on multiple genes in selective and independent manner or though coordinated interactions with each other is not clear.

**Fig. 1.**
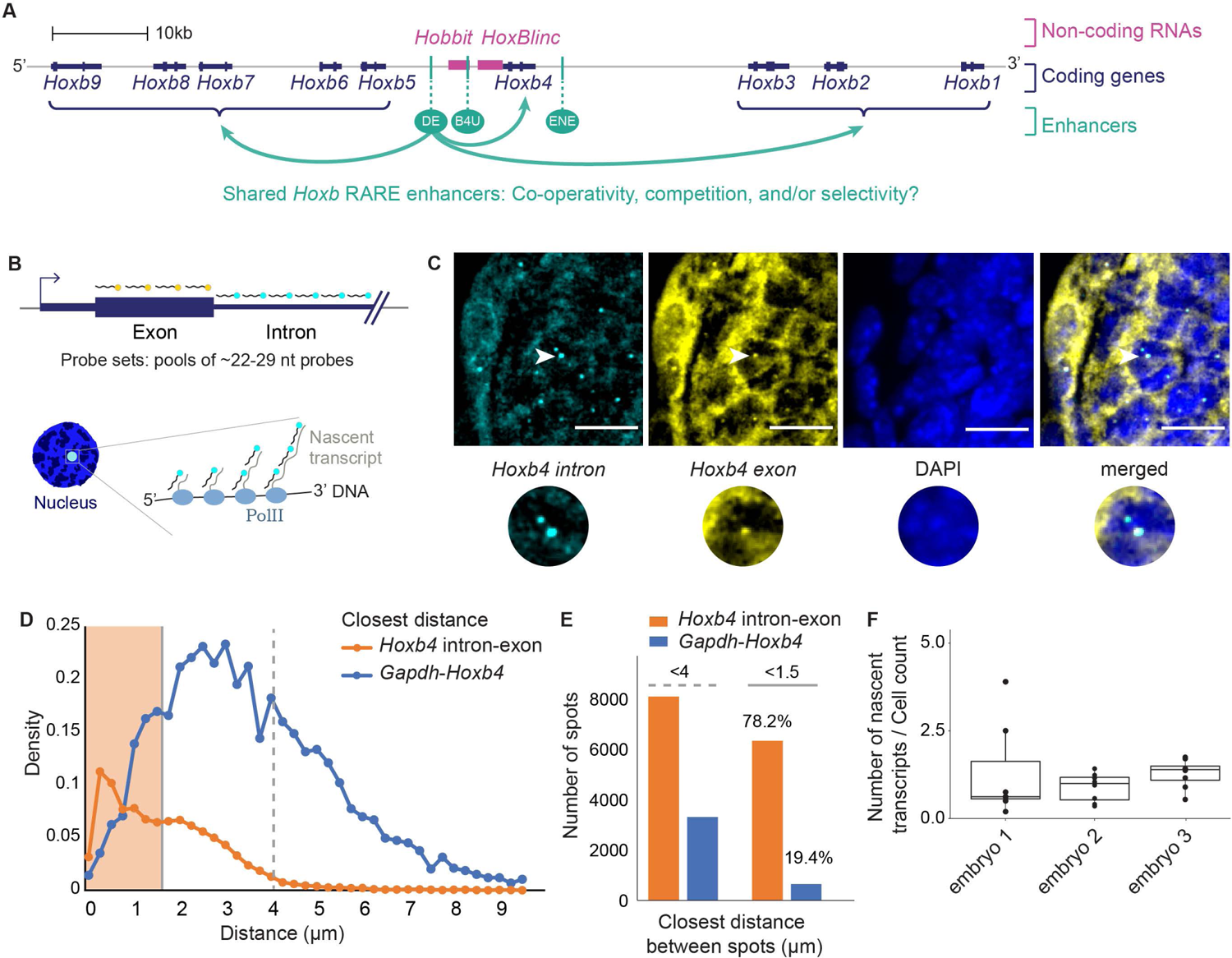
Optimized smFISH technique for detecting nascent transcripts at the level of single cells in mouse embryonic tissues. **A** Schematic of the *Hoxb* cluster with coding genes, non-coding RNAs, and RARE enhancers, drawn to scale. Previous evidence has shown that DE (arrows drawn) and ENE are able to globally regulate the steady state levels of genes in the cluster. **B** Probes design against gene introns and exons to detect nascent transcripts. Multiple nascent transcripts are made at regions of active transcription by many RNA Pol-II’s that are present. Upon hybridization these nascent RNAs get bound to probes, and each site of active transcription is visible as one nascent transcript in the nucleus of a cell. **C** Image of *Hoxb4* intron and exon probes. Arrow points at nascent transcripts that are shown also with a zoomed inset. Scale bar = 10 μm **D** Plot for closest distance between nascent transcripts from *Hoxb4*-intron and exon probes compared against *Hoxb4-Gapdh* probes. **E** Histograms drawn for closest distance spots with two thresholds: 4 μm for nuclear distance and 1.5 μm for co-localization. **F** Average *Hoxb4* nascent spots in tail sections of multiple embryos.

To begin to address these questions, we analyzed patterns of *de novo* (nascent) transcription of multiple *Hoxb* transcription units at the level of individual cells in mouse embryos. We employed a single molecule florescent *in situ* hybridization (smFISH) technique (Raj and Tyagi, 2010) in conjunction with probes spanning introns, and optimized it for use in tissue sections from stage matched wild type and mutant mouse embryos. Visualizing nascent transcripts in fixed tissue sections enabled us to obtain transcriptional snapshots at a defined stage in specific tissues to assess the *in vivo* transcriptional activity state of both coding and non-coding genes within the cluster. To quantify imaging data for a series of near adjacent whole tissue sections in an unbiased and high-throughput manner, we developed a deep learning (DL) algorithm for automatic calling and localization of signals for nascent transcripts in the nucleus. With the use of multiplexed probe sets, this imaging and DL pipeline allowed us to systematically quantify and compare patterns of nascent transcripts within the *Hoxb* cluster and their co-occurrence in individual cells. To assess the relative functional contributions of the three shared RA-dependent enhancers (*DE*, *B4U* and *ENE*) on *Hoxb* transcription *in vivo*, we generated mouse lines carrying a series of single and compound mutations in their RAREs and monitored the impact of these mutations on patterns of nascent transcription. Our analyses focused on neural tissue in the tail region of both wild type and RARE mutant embryos, where it is established that there are overlapping domains of expression of all *Hoxb* genes. We see no evidence for co-transcriptional coupling of all genes in the cluster in an individual cell. In wild type embryos, we predominately detect nascent transcription of a single gene in each cell, with a much lower frequency of simultaneous transcription of one or two other transcriptional units. Analyses in RARE mutant embryos reveals that individually each of the three enhancers differentially impacts patterns of nascent transcription of both coding and non-coding *Hoxb* transcripts. Two enhancers have a more global impact on transcription, while the third plays a predominantly local role impacting nascent transcription patterns in the cluster. The results suggest that selectivity and competitive interactions between enhancers plays a role in coordinating patterns of nascent transcription of *Hox* genes in the neural tube. This study provides new insights into the properties and *in vivo* roles of shared enhancers in coordinating the transcriptional dynamics of genes during mouse development.

## Results

### Developing an *in vivo* approach to study transcriptional regulation of the *Hoxb* cluster by shared enhancers

*Hoxb* genes are direct transcriptional targets of graded cues from RA signaling through a series of RARE-dependent enhancers embedded within and flanking the cluster (Nolte et al., 2019). The focus of this study is to investigate the properties of three shared RA-dependent enhancers (*DE*, *B4U*, and *ENE*) present in *Hoxb* cluster (Fig. 1A) and their input in regulating patterns of nascent transcription *in vivo* at the level of single cells in mouse embryos. The high density of enhancers and transcription units in the *Hoxb* cluster raises questions regarding promoter target selection, the degree to which these three enhancers individually or collectively participate in modulating short and long-range regulation of multiple genes and whether there is co-transcriptional coupling of regulated transcriptional units.

To begin to address these questions, we wanted to analyze patterns and sites of active transcription, which can be visualized by detecting nascent transcripts at the level of individual cells. Towards this goal, we employed a smFISH technique (Raj and Tyagi, 2010) and optimized parameters to facilitate its use in tissue sections from stage matched wild type and mutant mouse embryos (Fig. 2A). We designed and validated probe sets spanning introns for genes of interest (Fig. 1B) which enabled us to visualize nascent nuclear transcripts *in vivo* in fixed tissue sections. This optimized technique allowed us to monitor and quantify patterns of nascent transcription to obtain transcriptional snapshots at a defined stage in specific tissues and assess the *in vivo* transcriptional activity state of both coding and non-coding genes within the cluster at the level of individual cells.

**Fig. 2.**
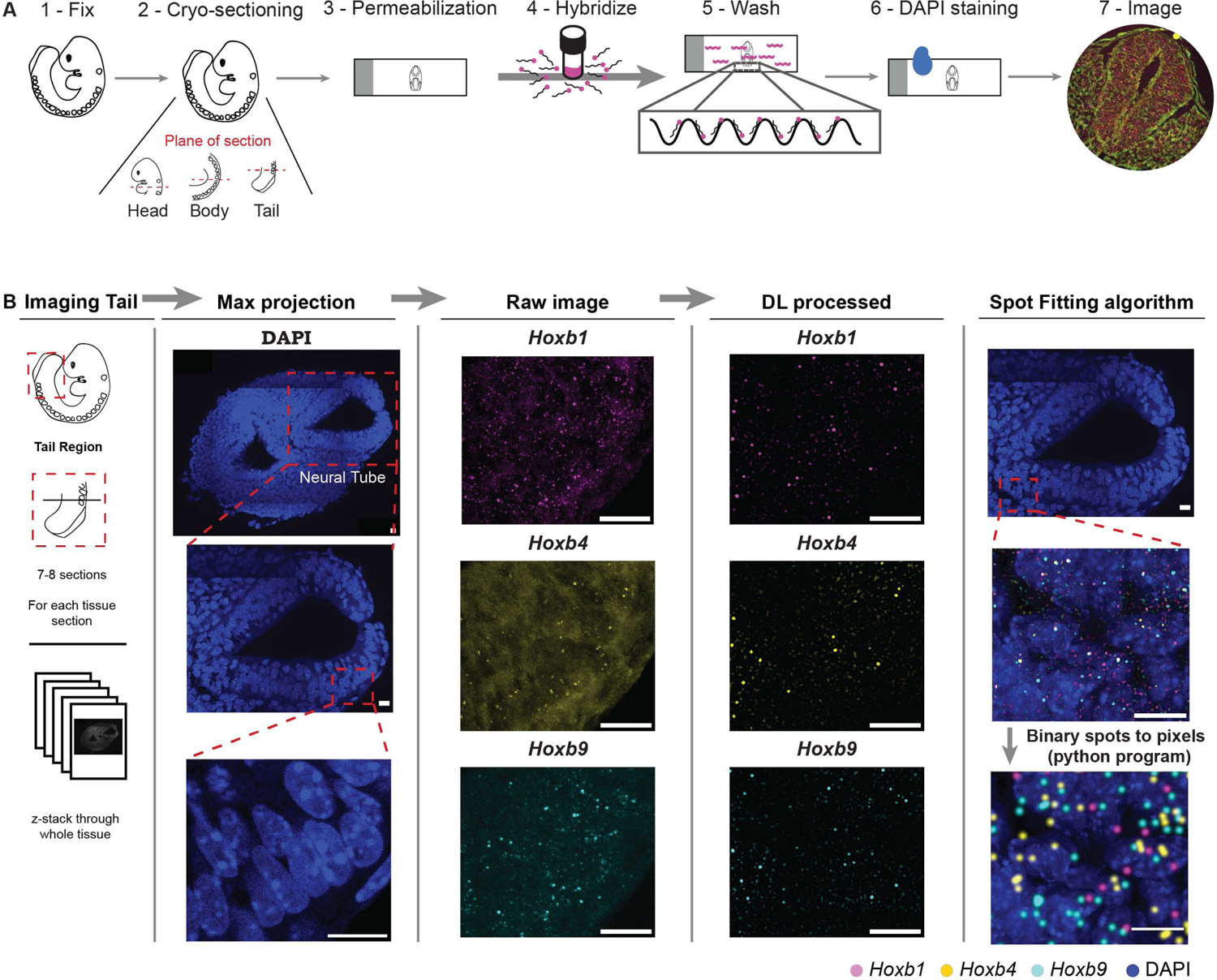
smFISH pipeline for processing tissue samples and quantifying nascent transripts. **A** Schematic for optimized single molecule Florescent in situ Hybridization (smFISH) technique, to detect nascent transcripts in cryo-sectioned mouse embryos. **B** Deep Learning (DL) pipeline to quantify nascent transcripts. A region of the neural tube is zoomed in to show raw image and the DL processed image. DL processed imaged can be spot fitted to increase the size of nascent spots for better visualization.

On each section, we employed multiplexed probe sets for up to 3 different transcripts and used DAPI staining to identify nuclei. Using alternate sections for different probe sets, we monitored and compared nascent transcription patterns of up to 6 transcription units within a defined region of the same embryo. Imaging through the whole tissue section (10 µm) allowed us to visualize all nascent transcripts present in the section. To circumvent the laborious task of manual counting, we developed a Deep Learning (DL) method to identify and quantify nascent transcripts in a high-throughput and unbiased manner (Fig. 2B). This imaging and DL pipeline allowed us to systematically quantify and compare patterns of nascent transcription of *Hoxb* genes and their co-occurrence in individual cells.

The DL output has a confidence probability between 0 and 1, for whether or not each detected fluorescent spot is a nascent transcript, over the whole tissue section. To test whether the network was performing well, we compared DL detected spots against spots marked manually and independently by several people on the same tissue sections. We found that there was an 80% overlap between different people marking nascent spots, and between people marking spots compared with those detected by DL on the same tissue sections (Fig. S1A). We interpret this as the DL network being as reliable as people in detecting nascent spots for a given section. For each tissue section, visual heatmaps for the intensities of nascent transcripts over the whole tissue were also generated (Fig. S2A). To better visualize nascent transcripts over the whole tissue, we performed spot fitting on DL detected nascent transcripts which allowed us to increase the size of the nascent spots and overlay them onto DAPI staining (Fig. S2B).

To verify the specificity of our probes in detecting nascent nuclear transcripts, we ensured the probes only showed 1 or 2 nuclear spots in a cell and have a high signal-to-background ratio (Fig. 1C). To further validate this approach, we hybridized the same tissue section with different probes sets for the *Hoxb4,* intron only and spanning the exon and intron, and observed a high degree of overlap in signals for nascent transcripts with both probe sets (Fig. 1C, zoomed insets). This shows that probe sets were indeed specific to *Hoxb4* and could robustly bind to *Hoxb4* nascent transcripts. The *Hoxb4* intron probe showed brighter localized nuclear fluorescent spots with minimal background, while the exon probe detected both nascent transcripts in the nucleus and weaker signals in the cytoplasm. We attribute this cytoplasmic signal as being indicative of single mature RNA transcripts but were unable to distinguish these transcripts from the autofluorescence background also present in the tissue. A caveat of our optimized smFISH protocol is that we could not distinguish single transcripts, hence we cannot measure total fluorescent intensity to infer transcriptional rates for genes (Fig. S1B). Therefore, in all smFISH analyses we focused on visualizing and quantifying only patterns of nascent transcription.

To further quantify the accuracy and robustness of our approach along with specificity of probes against nascent transcripts, we calculated the overlap of *Hoxb4* intron and exon probes. We compared distances between nascent transcripts observed from *Hoxb4* intron and exon probes against the distance between nascent transcripts observed for the *Hoxb4* exon and *Gapdh* control probes. In tissue sections, we cannot demarcate cell and nuclear boundaries, and hence measured nearest neighbor distances for all nascent spots detected in the whole tissue (Fig. 1D). As each nucleus is roughly 4 µm in diameter, we used that as a cut-off to obtain a count for all spots that are present within a single nucleus. For co-localization of spots from the *Hoxb4* intron and exon probes sets the peak distance was 0.3 µm, and the average distance between these spots was 1.87 µm (Fig. 1D). In contrast, for the *Gapdh* and *Hoxb4* spots the peak and average distances were 2.92 µm and 3.73 µm, respectively. To calculate the percentage of co-localized spots within cells, we divided the number of spots at distances <1.5 µm (co-localized or overlapping sites of nascent transcripts) by the number of spots within the same nucleus (<4 µm) (Fig. 1E). As expected for probes against the same transcript, we see that *Hoxb4* intron and exon probes overlap at a much higher frequency with smaller distances than the *Gapdh-Hoxb4* probes (Fig. 1D, E). Visual inspection of *Hoxb4* intron-exon spots appears to show that nearly all spots overlap. However, we observed an overlapping percentage of 78.2%, which may be attributed to co-transcriptional splicing. In this case, while we would capture spliced nascent transcripts with the exon probe, these would not be detected by the intron probe set. With respect to the *Gapdh* and *Hoxb4* probes, as expected the overlaps in signal were low (19.4%) (Fig. 1E). The actual overlap of these probes within cells is likely to be even lower because some of the scored *Gapdh-Hoxb4* spots <1.5 µm might also include instances where nascent transcripts from each gene are at the periphery of two different cells instead of being in the same cell.

To facilitate consistency in measurement of nascent transcripts, all 9.5 dpc mouse embryos selected for smFISH analyses were designated as being at the same specific stage based on their having an identical number of somites (24 somites). Sectioning and imaging settings were also kept consistent to ensure robustness of the technique. To examine consistency between embryos we quantified and compared the numbers of nascent *Hoxb4* transcripts in the tail sections of the neural tube from three different embryos (Fig. 1F). While there are some small differences in numbers of nascents observed in the individual tail sections, the averages are not significantly different between the three sets of sections. This implies that we can robustly quantify expression of multiple nascent transcripts and use averages of sections in a defined axial region of the tail for comparisons across different embryos.

### Patterns of nascent *Hoxb* transcription in wild type embryos

We next applied this approach to compare patterns of nascent transcription of multiple transcription units in the *Hoxb* cluster by designing and validating probe sets for coding and non-coding RNAs (Fig. 3A, Table S1). In addition to *Hoxb4*, we successfully generated robust probe sets effective for detecting nascent transcripts of the *Hoxb1* and *Hoxb9* coding genes at either end of the cluster, and for the lncRNAs, *Hobbit* and *HoxBlinc,* present in the center of the cluster. We attempted to generate probes for several other *Hoxb* genes, but encountered issues with high backgrounds, specificity, and reproducibility. Hence, we focused our analyses using the robust and specific probes sets we successfully developed.

**Fig. 3.**
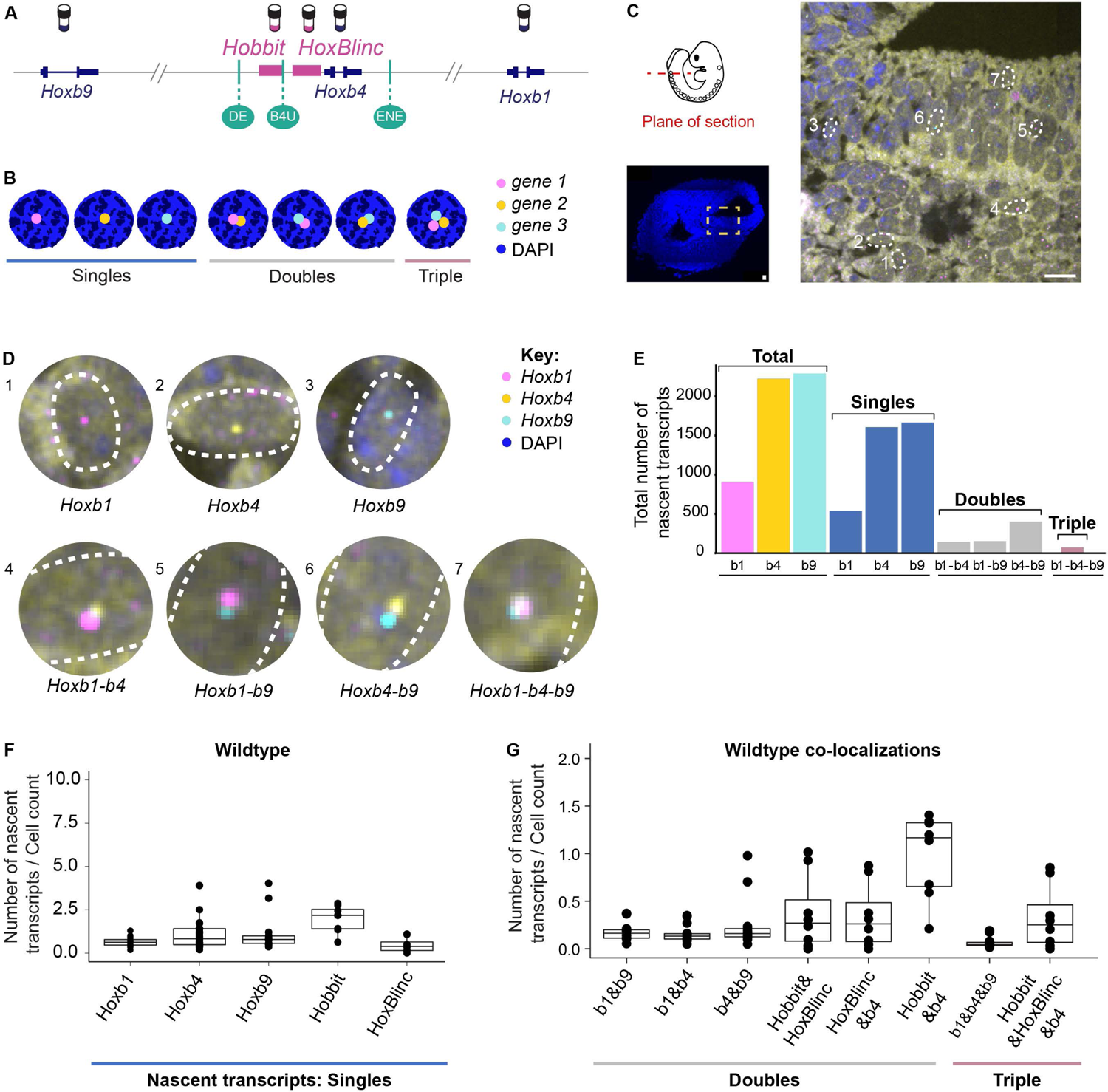
Quantifying patterns of nascent transcription of coding and non-coding genes from the *Hoxb* complex in wild type embryos. **A** smFISH probes designed against the ends and center of the *Hoxb* cluster. **B** Schematic for possible combinations of nascent transcripts that can be observed when hybridizing with three genes simultaneously. **C** Cross-section through one tissue sample from the tail region of a 9.5 dpc mouse embryo. Zoomed in region shows nascent transcripts present in portion of the neural tube. Scale bar = 10 μm. **D** Amplified cell insets from the image in C with spots marking nascent transcripts of *Hoxb* coding genes, showing data indicating examples of single, double, and triple co-localized transcripts. **E** Bar plot comparing the total number of nascent *Hoxb* transcripts measured for each gene in the single tail section shown in C & D. **F** Box plot comparing the total number of nascents/cell for *Hoxb* transcripts of the coding and non-coding genes calculated as an average of data from multiple near adjacent tail sections (7-9) of the same embryo. **G** Box plot showing the total number of co-localized double and triple combinations of nascent/cell for *Hoxb* the coding and non-coding genes calculated as an average of data from multiple near adjacent tail sections (7-9) of the same embryo. Each box in F and G represents the 75th (top line) and 25th (bottom line) percentile while the middle line represents the median expression.

Two different probe set combinations were used on a series of alternating tissue sections to examine the patterns of nascent transcription for coding and non-coding transcripts in directly comparable regions across the whole embryo. Probe set1 consisted of *Hoxb1*, *Hoxb4*, and *Hoxb9*, while probe set2 consisted of *Hobbit*, *HoxBlinc*, and *Hoxb4*. *Hoxb4* served as an internal control in both probe sets to verify that we detected comparable expression patterns when quantifying nascent transcripts across alternate sections (Fig. S3A). This allowed us to quantify nascent transcripts for 5 different transcription units in the posterior neural tube of each mouse embryo. We observed some minor variation between biological samples of embryos and in expression along the A-P axis of an embryo, but in wild type embryos the patterns compare very well when analyses are accurately conducted at the same axial level and planes across samples (Fig. S3B). As another means of evaluating the accuracy and specificity of this method, we quantified the patterns of expression of nascent transcripts of the 3 *Hoxb* coding genes along the A-P axis in wild type embryos by generating sections from the head, trunk, and tail regions (Fig. S4). We then compared the patterns of nascent transcripts along the A-P axis for these genes against the known profiles of their steady state expression determined by conventional colorimetric *in situ* analyses (Goh et al., 1997; Maconochie et al., 1997; Gould et al., 1998; Folberg et al., 1999; Brend et al., 2003; Medina-Martinez and Ramirez-Solis, 2003) and found they were in good agreement with the expected spatial order of expression along the A-P axis.

We performed analyses on the neural tube in the tail region where all the *Hoxb* coding genes and lncRNAs are known to be expressed and regulated by RA signaling. In each of the two multiplexed probe sets we simultaneously imaged up to 3 different transcripts. We observed a combination of signals which correspond to the activity of only one of the genes (singles) or co-localizations of nascent transcripts for 2 or all 3 genes, which we refer to respectively as doubles and triples (Fig. 3B). By imaging z-slices through a tail section, it is possible to visualize single and co-localized spots of nascent transcripts for the 3 coding genes (Fig. 3C). Magnified insets, illustrate examples of all of the combinations of co-localization of nascent transcription observed from the tail section (Fig. 3D). All of the nascent spots detected in the neural tube by the DL pipeline were quantified and plotted for each section, and the total number of signals for each of the nascent transcripts sub-divided into the categories of single, doubles, or triples based on their patterns of co-localization (Fig. 3E). For each probe set, ∼7-9 tissue sections from a tail region were imaged, and an estimate of total number of cells present in each neural tube was calculated for each section (see methods). The number of nascent transcripts were then divided by the cell count, to obtain the average number of nascent transcripts per cell for all 5 transcription units from analyses of both probe sets (Fig. 3F). We also determined the average number of co-localized double and triple categories of nascent transcripts per cell from all sections (Fig. 3G). Analyses of the patterns of co-localized nascent transcripts from a single tail section (Fig. 3E) or from the average of multiple sections (Fig. 3G) clearly show that we predominately detect nascent transcription of a single gene in each cell, with a much lower frequency of simultaneous transcription of one or two other transcriptional units.

These results in wild type embryos indicate that robust co-transcriptional coupling of all genes in the *Hoxb* cluster (coding and non-coding) is not the predominant mode of their coordinate regulation in individual cells, suggesting a rapid and dynamic process for potentiating transcription of individual transcription units. The low numbers of co-localized double or triple transcripts in wild type embryos indicate that, even in regions of overlapping expression domains in the tail region of the neural tube, nascent transcription of *Hoxb* genes is not simultaneously activated in the same cell at the same time. The data show that *Hobbit* and *Hoxb4* transcripts have a higher degree of co-localization than *HoxBlinc* and *Hoxb4* (Fig. 3G). If the non-coding RNAs, *Hobbit* and *HoxBlinc*, were run through transcripts of the transcriptional machinery activating genes sequentially along the cluster, then the *HoxBlinc* and *Hoxb4* would be expected to have similar co-localization patterns as *Hobbit* and *Hoxb4*. The differences we observed in the patterns for these lncRNAs suggest that transcriptional activation of these non-coding RNAs within the cluster is independently potentiated by distinct inputs.

### Assessing the role of shared enhancers using embryos with RARE mutations

Having characterized the patterns of nascent transcription of *Hoxb* genes in wild type embryos, we next sought to assess the relative functional contributions of the three shared RA-dependent enhancers (*DE*, *B4U* and *ENE*) on *Hoxb* transcription *in vivo*. For this goal, we generated mouse lines carrying a series of single and compound mutations in their RAREs and monitored the impact of these mutations on patterns of nascent transcription. Using CRISPR/cas9 gene editing approaches we generated a series of specific mutations in the *DE-*, *B4U-*, and *ENE-RAREs* by making base pair substitutions known to disrupt RAR/RXR binding sites (Sucov et al., 1990; Umesono et al., 1991; Oosterveen et al., 2003b; Ahn et al., 2014; Qian et al., 2018), but maintain the same spatial distances between these and other *cis*-elements in the endogenous genomic locus (Fig. 4A and Table S2). For each single RARE mutant, changes in the average number of nascent transcripts per cell count were visualized in the neural tubes of comparable tissue sections across stage matched wild type and RARE mutant embryos. We applied the spot fitting algorithm to aid visualization of the changes occurring in patterns of nascent transcription. Fig. 4B provides an example of the data and changes observed for nascent transcripts of *Hoxb1* in each of the three single RARE mutants compared to wild type, and results for all 5 coding and non-coding transcripts are presented in Fig. S5A-D. To facilitate comparisons of the datasets quantifying the number of nascent transcripts per cell for all 5 transcripts in the different genetic backgrounds, the results were plotted by grouping analyses according to each gene (Fig. 4C) or each individual RARE mutant (Fig. 4D).

**Fig. 4.**
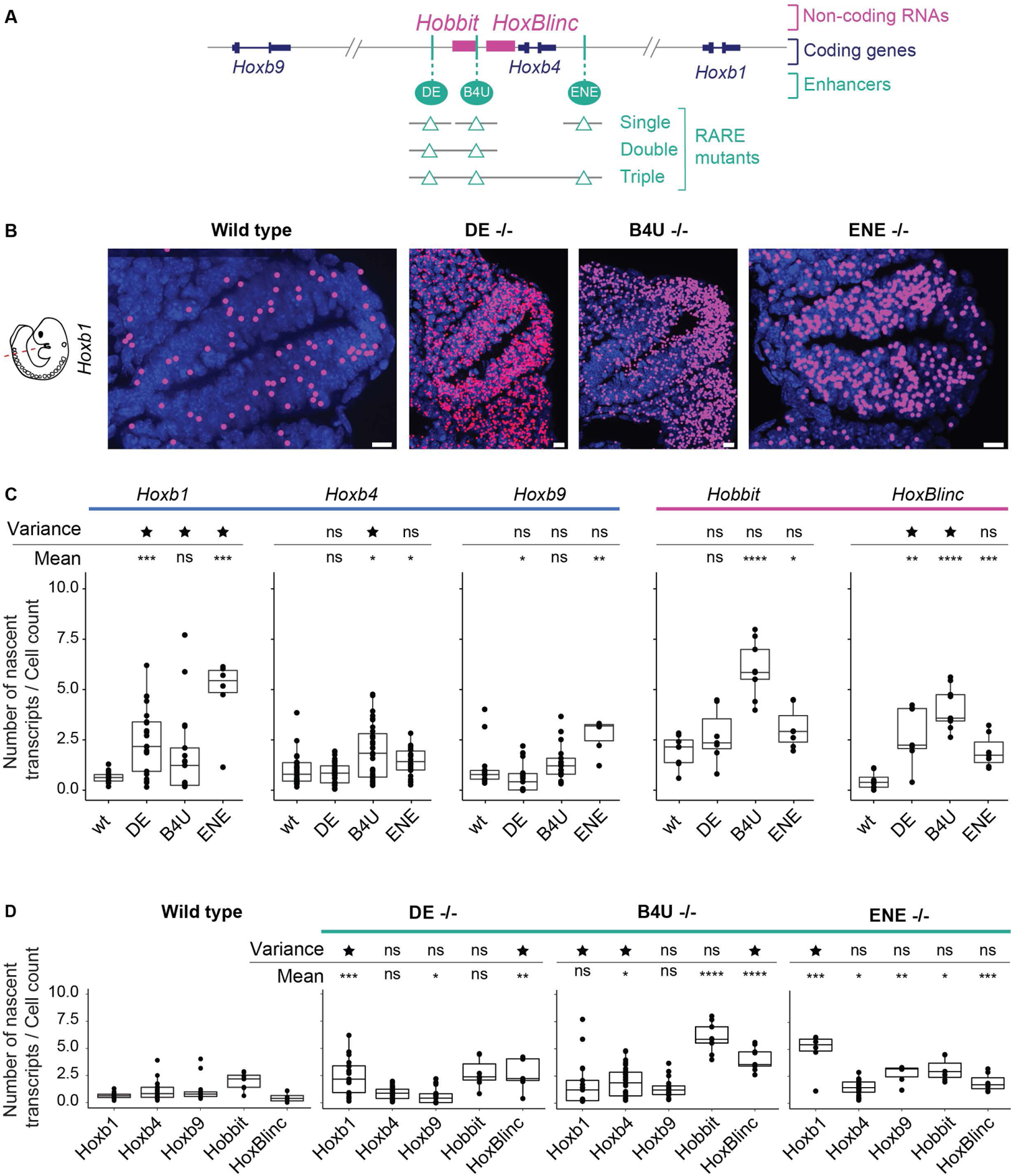
Quantification of changes in levels and patterns of nascent transcription of *Hoxb* coding and non-coding genes in single mutants of three different shared RARE enhancers. **A** Diagram depicting a series of single and compound mutants generated in RAREs of three shared enhancers present in the center of the *Hoxb* cluster. **B** Images of nascent transcripts for *Hoxb1* in wild type and the series of single RARE mutant embryos. The nascent transcripts detected by DL were spot fitted to increase the size of the spots for better visualization over the neural tube. **C** Box plot showing changes in average levels of nascent expression of coding and non-coding transcripts measured with two different probe sets on a series alternate tissue sections from multiple embryos of wild type and mutant mouse lines. The plots are grouped according to results for each gene. **D** Box plot of data from the same samples as in C, but grouped according to results for each genetic background. Each box in C and D represents the 75th and 25th percentile while the middle line represents the median expression. Significance is calculated based on both the mean (to estimate how the average expression is changing) as well as for the variance (to estimate how the range of expression among sections is changing) compared to wild type embryo sections. The mean differences that are significant compared to wild type are denoted by asterisks, while the variance values that are significant are marked by a star.

This analysis showed that levels of nascent transcription of *Hoxb1, Hoxb9* and *HoxBlinc* were altered in *DE*-*RARE* mutants, with elevated levels for *Hoxb1* and *HoxBlinc,* while *Hoxb9* displayed a slight decrease (Fig. 4C, D). In *ENE-RARE* mutants, *Hoxb1*, *Hoxb4*, *Hoxb9* and *Hobbit* all displayed levels of increased nascent transcription. In *B4U-RARE* mutant embryos, levels of *Hoxb4*, *Hobbit* and *HoxBlinc* all displayed an increase over wild type. These diverse changes in the mutants imply that all three enhancers have distinct inputs or preferences in regulating both coding and non-coding *Hoxb* transcripts. Furthermore, they indicate that the *DE-* and *ENE-RAREs* appear to have a greater long-range role in coordinately regulating transcriptional units spread throughout the cluster, while the *B4U*-*RARE* appears to be acting more locally in modulating *Hoxb4* and the *Hobbit* and *HoxBlinc* lncRNAs. These diverse regulatory changes in co-regulation of *Hoxb* genes in the mutants are also observed in evaluating patterns of co-localization of double and triple nascent transcripts (Fig. S6). Co-localizations of *Hoxb1*-*Hoxb4* are most impacted in the *DE* and *ENE* mutants, while *Hoxb1*-*Hoxb9* and *Hoxb4-Hoxb9* co-localizations are most impacted in the *ENE* mutant. Hence, mutating the *DE-* and *ENE-RAREs* impacts co-localization patterns of coding genes more than the lncRNAs, and conversely mutation of the *B4U-RARE* changes co-localization patterns for *Hobbit* and *HoxBlinc*, and the coding *Hoxb4* gene.

We wondered whether the number of co-localized nascent transcripts for multiple genes in wild type and mutant embryos was simply an outcome of random chance. Based on the number of single nascent transcripts for each transcription unit, we calculated the probability of random chance for co-localized transcripts. This analysis revealed that the actual observed co-localization trends were lower than what one would expect by random chance (Fig. S7). This suggests the presence of layers of regulation and enhancer interactions governing patterns of nascent *Hoxb* expression may include some that restrict or limit transcriptional co-activation.

### Coordinated transcription of *Hoxb* genes disrupted in compound RARE mutants

It was surprising that the general trend in changes to nascent transcription observed in the single RARE mutants reflected increased levels of transcription of selected subsets of genes rather than decreased activity. This suggests that the single RARE mutants may change the normal balance of regulatory interactions in the cluster, allowing the remaining functional RARE-dependent enhancers to shift or alter their target preferences resulting in functional compensation and changes in patterns of transcriptional activation of *Hoxb* genes. To investigate the degree to which these 3 shared enhancers can compensate, antagonize, compete, or work together to ensure coordinated transcription, we generated and analyzed compound mutants carrying the same alterations in each RARE.

Analysis of the *DE-B4U* double mutant, revealed that the elevated number of nascent transcripts per cell for *Hobbit* and *HoxBlinc* observed in the single RARE mutants unexpectedly returned to near wild type levels (Fig. 4C, 5A). Levels of nascent transcription of *Hoxb1*, *Hoxb4*, and *Hoxb9* coding genes were also more similar to that of wild type embryos. This suggests that there is antagonism or competition between the *DE* and *B4U* enhancers which is altered in each single mutant but eliminated in the double mutant. The fact that the nascent transcript levels trend towards wild type in the double mutant, implies that the *ENE*-*RARE* enhancer may functionally compensate for the mutations in the *DE* and *B4U* enhancers to maintain expression, consistent with its global effects on transcription in analysis of single *ENE-RARE* mutants (Fig. 4C, D).

**Fig. 5.**
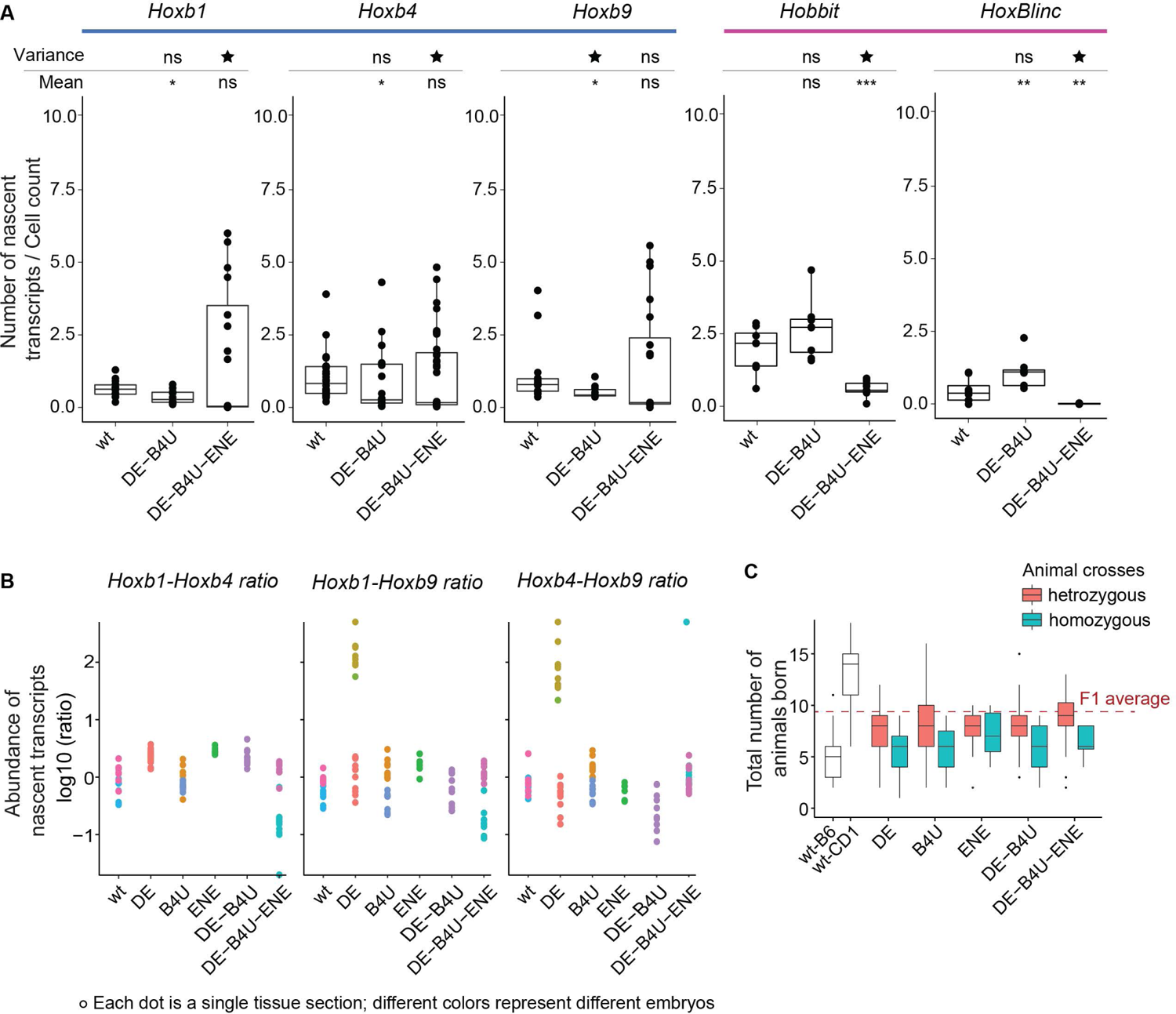
RARE mutations alter levels and proper coordinated transcription of *Hoxb* genes. **A.** Box plot showing changes in average levels of nascent expression of coding and non-coding *Hoxb* transcripts measured with two different probe sets on a series of alternate tissue sections from multiple embryos of wild type and compound RARE mutant mouse lines. The plots are grouped according to results for each gene. Each box represents the 75th and 25th percentile while the middle line represents the median expression. Significance is calculated based on both the mean (to estimate how the average expression is changing) as well as for the variance (to estimate how the range of expression among sections is changing) compared to wild type embryo sections. The mean differences that are significant compared to wild type are denoted by asterisks, while the variance values that are significant are marked by a star. **B** Abundance ratios of nascent transcripts/ cell plotted for the *Hoxb1*, *Hoxb4*, and *Hoxb9* coding genes. Each dot represents the ratio of counts from the neural tube for one gene over the counts for the second gene in the same section. All sections are for the tail region, and each color is a different embryo. **C** Litter sizes of the single and compound mutants, red for heterozygous animals and blue for homozygous animals. The two wild type strains from which F1 strains are generated for injections to make RARE mutants are shown in white. Dotted line shows approximate litter sizes for the F1 wild type animals.

To test this idea and investigate whether all three RAREs are essential for regulating *Hoxb* nascent transcripts, we generated and analyzed *DE*-*B4U*-*ENE* triple mutant embryos (Fig. 5A). We detected changes and considerable variability in the coordination of nascent transcription of genes in the *Hoxb* cluster in these triple mutants, with variable penetrance in the regulatory phenotype. For example, one mutant embryo displayed significantly reduced levels of nascent transcripts for all 5 genes, below levels in wild type, consistent with the 3 enhancers being required for potentiating activity (Fig. 5A). However, in another embryo, the levels and range of expression of nascent transcripts of the *Hoxb* coding genes in the tail was equal to or greater than wild type, but the levels for the *Hobbit* and *HoxBlinc* lncRNAs remained much lower than wild type. This extensive range of variation in levels of nascent transcripts for coding genes in the triple mutants, was generally not observed in our analysis of the single and double mutants.

However, we did observe an increase in the range of expression of *Hoxb1* and *HoxBlinc* in single *DE* and *B4U* mutants (Fig. 4C, D). To quantify changes in the range of levels of nascent transcription in the RARE mutants, we calculated whether the variance was significantly different in the series of mutants analyzed compared to wild type (Fig. S8). The significant changes in variance we detected in mutant embryos suggests that functional *DE-*, *B4U-*, and *ENE-RAREs* are important in coordinating and maintaining the proper levels of coding and non-coding nascent transcripts in the tail region.

We examined whether changes in variance were correlated between different *Hoxb* genes and plotted ratios of average nascent transcript levels of one coding gene over a second coding gene (Fig. 5B). We found that the relative levels of coding genes were very close to those of wild type in most RARE mutants. However, in the *DE-RARE* single mutant biphasic patterns of variation were observed for ratios of *Hoxb1*-*Hoxb9* and *Hoxb4*-*Hoxb9,* indicated by the presence of two populations of abundance ratios for the coding genes in these mutants (Fig. 5B). Similarly, in the *DE-B4U-ENE* triple mutant ratios of *Hoxb1*-*Hoxb4* and *Hoxb1*-*Hoxb9* displayed two distinct patterns. In *DE* mutant, these two populations can be attributed to the increased variance in *Hoxb1* nascent transcription (Fig. 4C), while in the triple mutant variations in both *Hoxb1* and *Hoxb9* contribute to the range of variability. These changes highlight that mutations in the RAREs can result in variable penetrance of their regulatory phenotypes in a manner that suggests they contribute to regulation of both the levels and robustness in patterns of nascent transcription.

The level of regulatory variability in the RARE mutants may be greater than we have detected because large changes and variability in coordinated *Hoxb* expression may affect development and viability of embryos. We have observed that mutations in the RAREs, result in compensatory mechanisms altering levels of *Hoxb* transcripts that are often greater than wild type (Fig. 4, 5) and there may be a limited range of thresholds in expression levels that are compatible with viability. Through variable penetrance, if mutations in the RAREs result in changes outside of this range these embryos may be lost. In generating crosses of the series of RAREs mutants to obtain embryos for analysis in this study we have noted an impact of the mutations on litter sizes (Fig. 5C). We get reduced litter sizes for most homozygous RARE mutants compared with their heterozygous littermates and all mutants have much lower numbers of viable animals born compared to the F1 wild type animals (Fig. 5C, dashed red line). In addition, homozygous animals seem to fare worse and indeed often have to be backcrossed to wild type animals to maintain viability of the mouse lines. This suggests that in homozygotes there may be some selective pressure, whereby the most severely affected mutant embryos are eliminated or resorbed. However, we are able to generate homozygous animals for all the RARE mutant lines. This indicates that while these RAREs are critical for proper coordinated expression of *Hoxb* genes, other regulatory elements must have inputs that contribute to levels of *Hoxb* transcription that allow viable development.

### Enhancer-promoter distances decrease in the RARE mutants

To explore what might be happening at the level of spatial relationships between genes in the RARE mutants, we examined the distances between co-localized (e.g. co-regulated) nascent transcripts in wild type and the single RARE mutants in cases where all three coding genes were transcriptionally active. For each sample, we identified the center of each nascent transcript spot and measured distances between the other co-localized spots (Fig. 6A). We then compared the distances between co-localized transcripts in RARE mutants versus wild type, to look for potential changes in the nature of enhancer-promoter interactions. We observed a range of distances between the nascent transcripts of co-localized spots in the wild type and mutant embryos (Fig. 6B) and plotted the peak distance between nascent spots (Fig. 6C). To visualize the changes in the distances between all three *Hoxb* coding transcript spots we also generated a triangle plot (Fig. 6D). As a control for each pair-wise distance comparison, we used *Hoxb4* intron-exon probe distances. This distance bar for *Hoxb4* intron-exon probes in wild type embryos acts as an indicator for our resolving power in distinguishing between two identical co-localized spots. For a negative control of co-localization, we utilized *Hoxb4-Gapdh* nascent spots. We found that there are not enough co-localized spots of *Hoxb4-Gapdh* to make a distance bar, as expected for genes that are not under the same transcriptional control and chromosomal locations.

**Fig. 6.**
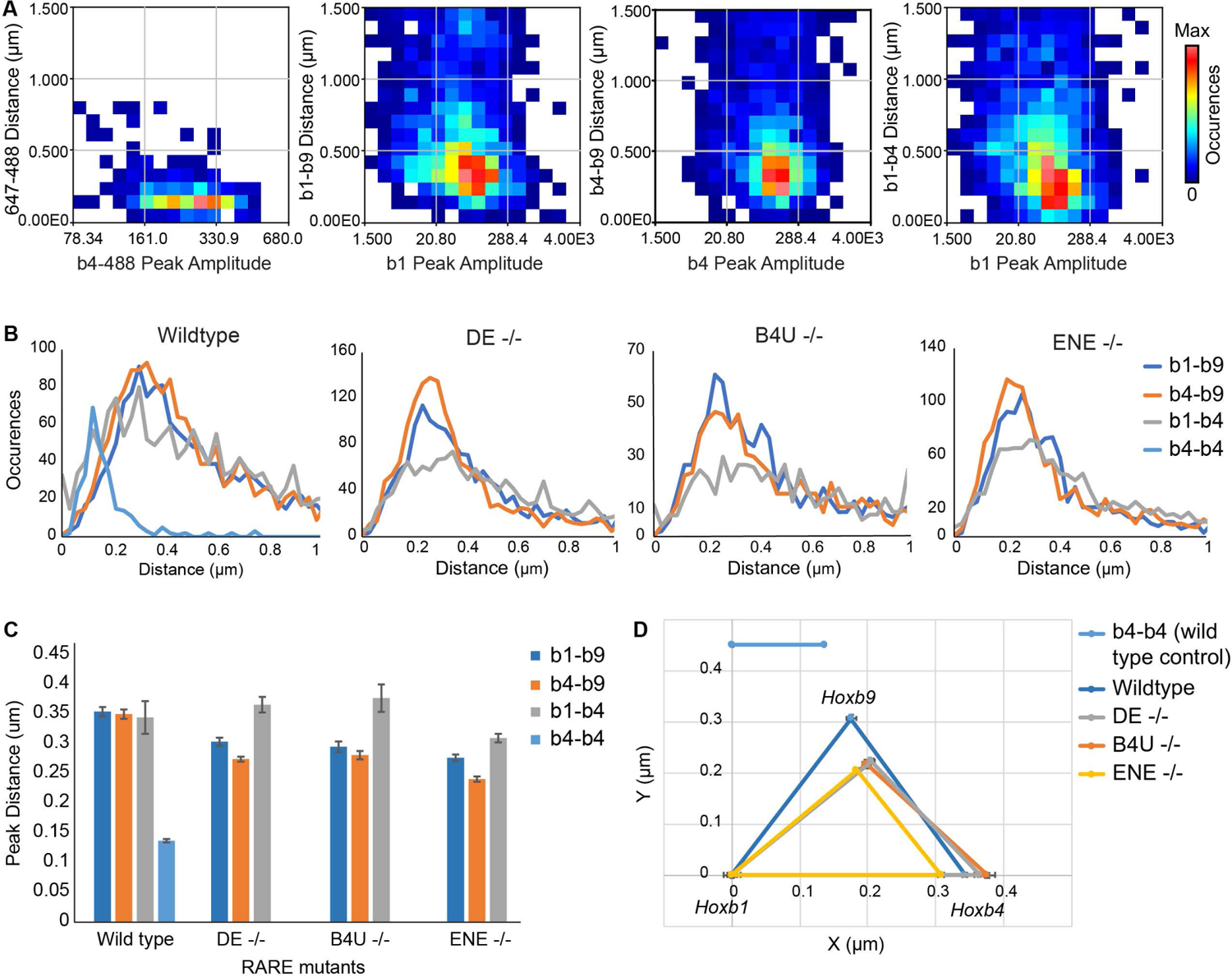
Distances between triple co-localized nascent transcripts of coding genes is decreased in RARE mutants. **A** Distances between nascent spots (y-axis) compared to a peak reference (x-axis). First panel on the left represents distances from data using control *Hoxb4* intron (647 probe) and intron plus exon (488 probe) probes; each subsequent adjacent panel represents distances between *Hoxb1-Hoxb9*, *Hoxb4-Hoxb9*, and *Hoxb1-Hoxb4*, respectively. The colors in the heat map indicate the frequency of occurrences of the nascent spots for corresponding distances. **B** Occurrence of nascent spots plotted against a gated distance of 0 to 1 μm. **C** Histogram for the peak values of the distance distributions between nascent spots. Colors of bars indicate which two gene spot distances are plotted, as indicated by key on the right. **D** Triangle plot for the peak distances observed between nascent spots of coding *Hoxb* genes in C. *Hoxb1* is anchored at 0, and relative distance changes between genes are plotted with respect to *Hoxb1*.

These analyses revealed that in wild type cells, the distance between the control probes for the *Hoxb4* intron and exon co-localized nascent spots is 136 nm (with a simulated standard error of 1.5 nm), setting the resolution threshold in our ability to distinguish distances between any two spots. *Hoxb1*-*Hoxb9,* which are ∼90 kb apart in the cluster, have nascent transcripts that are separated by a peak distance of 350 nm (Fig. 6C, D). This distance is similar to that previously observed for co-regulated genes at comparable genomic distances in other systems (Chen et al., 2018; Benabdallah et al., 2019; Levo et al., 2022). *Hoxb1*-*Hoxb4* and *Hoxb4*-*Hoxb9* which are ∼44 kb and ∼42 kb apart, respectively, are also separated by a mean distance of 350 nm (Fig. 6C, D). In the single RARE mutants, we observed that the distances between *Hoxb1*-*Hoxb9* and between *Hoxb4*-*Hoxb9* decreased, while there was no significant difference in the *Hoxb1*-*Hoxb4* distance. The finding that *Hoxb9* nascents appear closer to *Hoxb1* and *Hoxb4* in each of the RARE mutants is consistent with the idea that a mutation in one enhancer may change the dynamics of regulatory interactions and spatial organization of the other enhancers in the cluster during co-transcriptional activation of multiple *Hoxb* genes.

## Discussion

In this study, we developed smFISH approaches with probes spanning introns to investigate how 3 shared RA-dependent enhancers embedded in the *Hoxb* cluster regulate patterns of nascent transcription of multiple genes *in vivo* at the level of single cells within the developing mouse neural tube. In wild type embryos we predominately detect nascent transcription of only a single *Hoxb* gene in each cell, with a much lower frequency of simultaneous transcription of one or two other transcriptional units. We find no evidence for simultaneous co-transcriptional coupling of all or specific subsets of genes in the cluster in individual cells. Analyses in embryos with single and/or compound mutations in RAREs of these enhancers revealed that each enhancer differentially impacts patterns of transcription within the cluster. The *DE* and *ENE* enhancers have a more global role in coordinating the levels and patterns of nascent transcription of multiple genes in the cluster, while the *B4U* enhancer plays more of a local role impacting nascent transcription of near adjacent genes. Changes we observed in compound mutants (summarized in Fig. 7) suggest that selectivity and competitive interactions between these enhancers plays an important role in coordinating and robustly maintaining the proper levels of nascent transcription of *Hoxb* coding and non-coding genes in the neural tube. Together these data suggest that rapid and dynamic regulatory interactions potentiate transcription of individual transcription units through combined inputs from these enhancers in coordinating the response to RA. This study provides new insight into the *in vivo* transcriptional dynamics of the *Hoxb* complex in mouse embryos and raises interesting questions about the mechanisms and the roles of shared regulatory elements in coordinating expression.

**Fig. 7.**
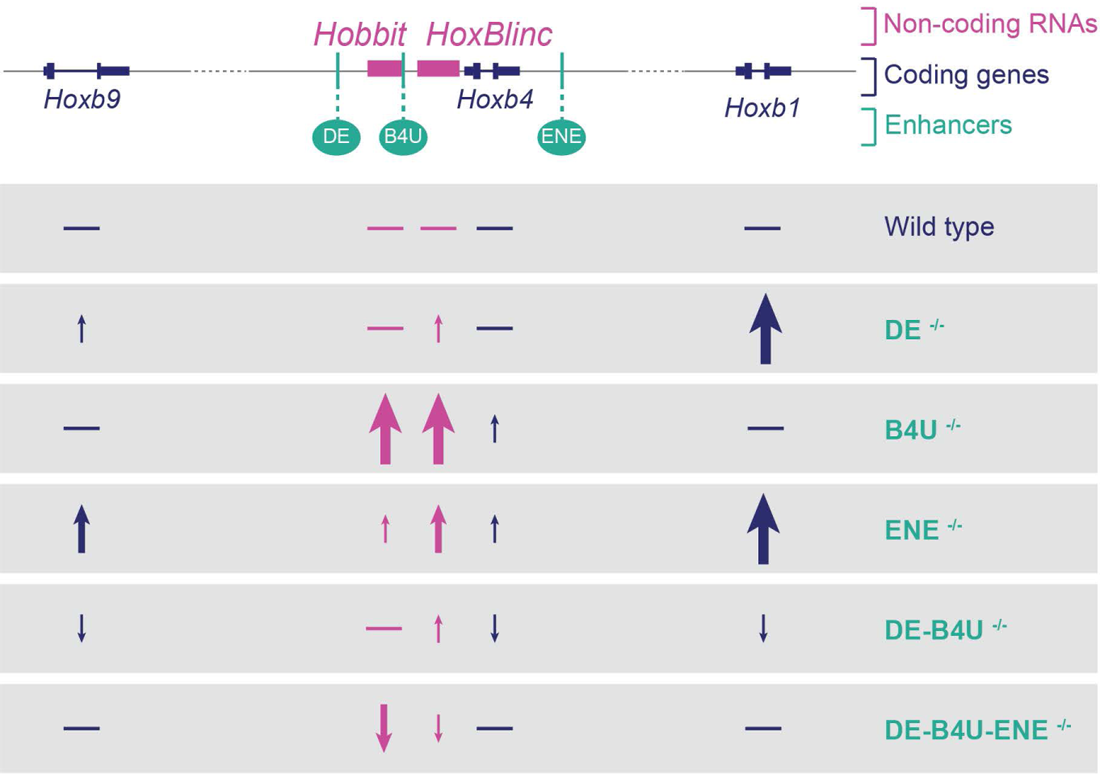
Summary of effects of single and compound RARE mutants on patterns of nascent transcription. Differences observed for coding and non-coding *Hoxb* genes in RARE mutants compared against wild type are depicted. Arrows indicate direction of change, increase (up) or decrease (down), relative to wild type expression. The thickness and height of arrow indicates the relative levels of change, with the thickest and tallest arrows indicating the most significant changes. Horizontal lines indicate levels similar to wild type.

In the smFISH analyses, we primarily observe single spots of nascent transcription for each of the transcriptional units and very low levels of co-localized transcriptional events for other genes. Transcription in eukaryotes is thought to occur in bursts that are spaced out by refractory periods (Bartman et al., 2016; Fukaya et al., 2016). Unfortunately, we are unable to infer transcriptional bursting rates because we cannot detect single transcripts, but the data suggests that at a given time only one allele is transcriptionally active. We do not find evidence for the equivalent of a ‘topological operon’ (Levo et al., 2022) that coregulates all of the *Hoxb* target genes through shared enhancers. We can’t exclude the possibility that there are sub-sets of genes in the cluster that are co-transcriptionally regulated, as we were unable to develop probe sets for all of the transcription units. While only one of the 5 genes we examined generally appears to be active at a given time, there is a low level of co-transcriptional activation of two or three other transcription units. This implies that there are likely to be rapid and dynamic enhancer-promoter interactions between each of the RA-dependent enhancers and their target genes during activation of transcription.

A surprising general finding in our analysis of single RARE mutants is that we observed increased levels of nascent transcription of selected subsets of genes (Fig. 4C, D), indicating a change in the normal balance of regulatory interactions in the cluster. This implies that altering the activity of each enhancer leads to an increase in the transcriptional bursting frequency or duration. Furthermore, in the *DE-B4U* double mutant elevated levels of nascent transcripts per cell seen in the *DE* and *B4U* single mutants unexpectedly returned to near wild type levels (Fig. 4C, 5A). This suggests that there is synergy, antagonism or competition between the *DE* and *B4U* enhancers which is altered in each single mutant but eliminated in the double mutant. This could arise through changes in enhancer-enhancer or enhancer-promoter interactions, or alterations in their individual or combined target preferences. The enhancers may also have optimal proximities between each other and their promoters to potentiate activity. Our analysis of co-localized nascents of *Hoxb1*, *Hoxb4* and *Hoxb9* in RARE single mutants revealed that *Hoxb9* nascents appear closer to *Hoxb1* and *Hoxb4* (Fig. 6). This is consistent with the idea that a mutation in one enhancer may change the dynamics of regulatory interactions and spatial organization of the other enhancers in the cluster during co-transcriptional activation of multiple *Hoxb* genes. These findings raise the possibility that the *Hoxb* genomic locus may have a 3D architecture which can serve as a transcriptional hub, in which the three RARE enhancers dynamically compete or synergize with each other in interacting with and activating their target *Hoxb* gene promoters. This could potentiate the rapid turning on or off of nascent transcription of individual, and sometimes multiple genes, in a controlled manner.

In analysis of wild type embryos, we noted that quantitation of levels of nascent transcripts for the *Hoxb* genes were very reproducible and showed little variation with a narrow range, even between embryos (Figs. 1F, 5A,B). However, in the single and compound RARE mutants there is a wide range of variation in levels of nascent transcripts for some but not all *Hoxb* genes. For example, in single *DE* and *B4U* mutants the range of expression of *Hoxb1* and *HoxBlinc* are much greater than wild type, while in triple mutants there is a biphasic pattern where levels can be well below or much greater than wild type (Figs. 4C, D, 5A, B). The significant changes in variance we observed in RARE mutant embryos suggests that functional *DE-*, *B4U-*, and *ENE-RAREs* are important for coordinating and maintaining the degree of variability and proper levels of *Hoxb* nascent transcripts in the tail region.

This study has highlighted the importance of these RAREs and shared enhancers in coordinating global levels of *Hoxb* expression in integrating the response to RA. While essential for proper expression of *Hoxb* genes, other regulatory elements and/or mechanisms contribute to regulating patterns of transcription of these genes. For example, in the triple RARE mutant embryos we find a wide range in levels of nascent transcripts, ranging from below to well above those of wild type. This suggests that mutations of the RARE may enable new or altered inputs from other regulatory regions of the cluster that compensate for the loss in activity of these shared enhancers. In conclusion, our findings provide new insight into the roles of shared regulatory elements in coordinating regulation of the *in vivo* transcriptional dynamics of the *Hoxb* complex in mouse embryos.

## MATERIALS AND METHODS

### Mouse lines

Wild type animals were generated from crosses of the Stowers F1 strain, which are a cross between CBA/CaJ x C57Bl/10J. The RARE enhancer mutants used for the project were generated by Christof Nolte and Youngwook Ahn in the Krumlauf lab using a CRISPR approach. The *DE* mutant was generated as previously described (Ahn et al., 2014). For generation of the *B4U* and *ENE* single mutants, *DE-B4U* and *DE-B4U-ENE* compound mutants, gRNA sequences (Table S3) were ordered as oligonucleotides (IDT) with adjacent cloning sites added and cloned into *pX330* plasmids following directions provided by the Zhang Laboratory (Cong et al., 2013). The replacement oligonucleotide (at 75 ng/µl) and the Cas9 targeting plasmid (at 3 ng/µl) were combined and microinjected into the pronuclei of 1-cell embryos. 2-cell embryos were transferred the following day into Stowers F1 recipient mice. All mutations introduced into the endogenous Hoxb locus were validated by sequencing to ensure that the specific base pair mutations were properly generated. All animals are maintained by Laboratory Animal Services (LAS) at Stowers Institute for Medical Research (SIMR) and all experiments involving mice were approved by the Institutional Animal Care and Use Committee of the Stowers Institute for Medical Research under a protocol (Protocol # 2021-134) issued to R. K.

### Genotyping of animals

To genotype the animals, tail samples were collected after weaning the animals. The tails were either processed internally and genotyped or directly sent for processing to Transnetyx. For each mutant animal, validated Transnetyx assays have been set up to detect the presence of wild type or mutant DNA sequences The link to the website is https://www.transnetyx.com/.

### Staging of embryos

For each embryo collected, the number of somites were counted to ensure same developmental age was used for all experiments. Embryos are considered 9.5dpc if they have anywhere from 21 to 29 somite pairs, but for this paper all wild type and mutant RAREs embryos containing 24 somite pairs were used for all experiments.

### Collection and fixing of embryo samples

For collecting wild type embryos, Stowers F1 animals were paired and pregnant females upon plug checking were marked for embryo collection at 9.5dpc. For enhancer mutants, mouse lines were bred to homozygosity. The homozygous animals were then crossed, and pregnant females upon plug checking were marked for embryo collection at 9.5dpc. The pregnant females were euthanized according to IACUC protocol number 2021-134 (detailed protocol is available at eprotocol.stowers.org). The uterine tissue containing embryo sacs were extracted from the abdominal cavity and kept in cold PBS solution. After collecting all 9.5dpc embryos in cold PBS, the embryos were transferred to 4% PFA solution and kept overnight at 4°C. The embryos were then dehydrated using methanol gradients of 25%, 50%, 75% and 100%. The embryos were stored in −20°C for at least one day before histological processing.

### Histological preparation of embryos

For smFISH RNAase free conditions were used. The embryos were kept in methanol and rehydrated slowly with PBS. The embryos were cut into three sections to aid with correct orientations; head (cut made after brachial arches), mid (cut made after hindlimb bud), and tail sections. After this the samples were dehydrated using a 15% solution of sucrose made in DEPC treated PBS. Once, the samples were saturated, they were embedded into OCT (VWR #25608-930) using the Histochill unit (SP Scientific Products # HC80). Using a Thermo Scientific Microm Cryostar NX70 cryostat, transverse sections at 10microns were placed on Sure Bond charged slides (Avantik #SL6332-1); Alternate sections were placed onto a single slide. The slides were stored at −80°C if not going to be used within the same week or kept at −20°C if going to be used within the same week.

### Probe design

For probe design genomic sequences were obtained from Genome Browser (genome.uscs.edu). The multiple 20-22 bp intronic and exonic probes (for *Hoxb4*, *Hobbit*, and *HoxBlinc*) and the multiple 29bp intronic probe (for *Hoxb9*) were designed using the Stellaris Probe generator from LGC Biosearch technologies (https://www.biosearchtech.com/support/tools/design-software/stellaris-probe-designer). The *Hoxb1* and *Gapdh* mouse probes were brought pre-labelled from the Stellaris website.

### Probe synthesis

For smFISH the probe-set sequences were either bought from the Design Ready version offered by Stellaris or were designed using the Stellaris probe designer tool. For probe labeling, unlabeled probe sets carrying a C-term TEG-Amino tag, were purchased from Biosearch Technologies and labelled as previously described (De Kumar et al., 2017). Probes (5 nmol) were fluorescently labeled overnight in 0.1 M sodium tetraborate pH 9 at 4 °C. Labeling occurred with AlexaFluor-488, AlexaFluor-568 or AlexaFluor-647. Two units of amine reactive succinimidyl ester Decapacks (ThermoFisher) were used for each reaction and following quenching labeled probes were purified using Reverse-Phase HPLC. Probes were separated using an Ettan LC (GE Healthcare) using a 4.6 × 250-mm, 5-μm, C18-EMS end-capped Kinetex column (Phenomenex). Mobile phase A was 0.1 M ammonium acetate (EMD) pH7 and mobile phase B was acetonitrile (Millipore). A linear gradient of 5% B to 100% (vol/vol) B was run over the course of 20 min at 1 mL/min. Peaks were monitored at 280 nm for probe and at 488, 568 or 647 nm, depending on the dye. Dual positive peaks were collected by hand and concentrated by spin vac. Samples were then resuspended in DEPC water to a final volume of 100µL. All probes and their fluorophores are in Table1.

### smFISH optimized protocol

The smFISH existing protocol (Raj and Tyagi, 2010) was optimized and adopted to use for mouse embryo sections. Tissue slides were thawed at room temperature for ∼10 minutes. The samples were permeabilized with 2ng/ml concentration of proteinase K in PBS for 10 minutes. Sample was kept in wash buffer A (Stellaris, Catalog # SMF-WA1-60), until hybridization mixture was prepared. 100ul of deionized Formamide (VWR Catalog # CAS-75-12-7) was added to 900ul of hybridization buffer (Stellaris, Catalog # SMF-HB1-10) to make the hyb solution. For every 100ul of hyb solution, 1ul of probe was added (∼5nmol stock probe dissolved in 100uL of nuclease free water), and up to 3 probes were added to each mixture and put on tissue slides. The slides were covered with a coverslip, heated at 65 °C for 10 minutes, and kept in a wet chamber overnight at 30 °C (∼16hours). Tissue slides were then washed with wash buffer A for 30mins at 37 °C. DAPI at a 1:5000 ratio was added to wash buffer A, and slides were kept in this mixture at room temperature for 10mins followed by 2-4 hours at 4 °C. Slides were washed with wash buffer B (Stellaris, Catalog # SMF-WB1-20) for 15mins at room temperature. Slides were mounted with prolong gold and covered with a 0.13-0.16mm long coverslip and kept to dry at room temperature before imaging.

### Imaging with Nikon spinning disk

Tissue sections on the slides were imaged using the Nikon Ti-E microscope coupled to a Yokogawa CSU-W1 spinning disk using the Hamamatsu Flash 4 sCMOS camera. The images were captured using 60X Plan Apochromat (NA 1.4) oil objective. The sections to image were manually identified using a 10X objective (NA 1.45). The images were obtained at 100% laser power for far red 633, red 561, green 488, and DAPI 405 nm lasers. For each channel the following filters were used: DAPI, ET455/50m, green, ET525/36m, red, ET605/70m, and far red, ET700/75m. A Nikon Elements Job “Tiler” was used to capture the images if tiles were taken, the image was stitched later in Fiji (https://imagej.net/software/fiji) using the grid/collection stitching plugin. The order of experiments was Lambda (z-series), so each color was imaged in z before moving to next color. For obtaining a z-stack through the tissue section, each slice imaged was 1 µm apart and a range of 20 steps was taken to be imaged. Images were obtained in the order of 633 (for the 647 probe), 561 (for the 555 probe), 488 (for the atto 488 probe), and 405 (for DAPI). On a slide the whole tissue section was imaged, and for each slide a single row was imaged all the way across for each genotype of embryo. All images were stitched using Fiji before further processing.

### Deep learning (DL) processing on images

RNAFish spots were segmented using DeepFiji (Nuckolls et al., 2020). In brief, a small subset of spots from different probes were manually annotated in Fiji and used to train a 2D Unet (Ronneberger et al., 2015) model. This model was used to infer spot probabilities from full image sets. After obtaining the results from DeepFiji, 3D image sets were thresholded, and 3D segmented in order to apply a size filter, and then projected to 2D. Combinations of these projections were examined to find cases where spots had overlap in multiple channels. Original spots and overlapped spots were then reduced to a single pixel and blurred to provide heat maps. Spot locations were also saved for future analysis. Relevant code for preparing image files for DeepFiji, and the post-processing afterwards can be found at https://github.com/jouyun/smc-macros.

### Spot Fitting

To be able to see nascent spots detected over the whole tissue section without having to zoom into image, the nascent spots localization file was used to make a spot file. To generate a spot file for this visual representation, apython program was used. The spot file could be edited in Fiji, such that the sizes of the individual spots could be increased. Spot size for each channel was increased consistently across all samples, and this file was overlaid onto DAPI.

### Distance measurement methods

Distances were measured between smFISH spots via Gaussian fitting with color correction utilizing custom plugins written for the open-source image processing program Fiji. These tools are available at https://research.stowers.org/imagejplugins. Color correction parameters were determined by fitting tetraspeck beads (Thermo Fisher) to Gaussian functions and finding the translation, rotation, and scaling parameters that minimize the distances between peaks in different channels. Those same transformations were applied to smFISH images collected with the same parameters. After color correction, images were pre-processed via Gaussian blurring with a standard deviation of 1 pixel and rolling ball background subtraction with a ball radius of 15 pixels. Peaks were then found in the far-red channel (due to it’s low background) by a max not mask (spot finding Fiji plugin) approach. Briefly, the maximum voxel in the image is repeatedly found and then a spheroid around that voxel is masked with zeros. For this analysis the spheroid diameter was 20 pixels in xy and 6 slices in z. This procedure is repeated until no other voxels are found with and intensity above 5% of the image maximum.

Once positions were found, they were refined by non-linear least squares 3D Gaussian fitting of 20 x 20 pixel stack of 6 slices surrounding the potential position in all 3 channels. Best fit peak positions were constrained to be within two pixels distance in xy and 2 slices in z of the maximum position found above. For each pair of probes, 2D histograms of spot amplitude vs. spot distance were created (see Fig. 6) and the area surrounding the main peak of this histogram extending from distances of 0 to distances of 1 micron was gated and plotted in D (Fig. 6). Finally, that histogram was fit to a one-dimensional Gaussian, again by non-linear least squares to obtain the peak position of the distance distribution.

### Measurements and analysis post-DL on images

Images were processed using Fiji. Using max projection from the DAPI channel, the regions of interest (ROIs) were marked in each tissue section. ROIs for the neural tube and adjacent somites were marked for each tail section by hand in the DAPI channel. Measurements were made for the total intensity of nascent transcripts for each ROI and from each ROI the background intensity was subtracted. For each ROI, a custom in-house written plugin “filter table spatial ROI jru v1” was used to also extract the exact count of nascent spots and their co-localizations from the DL processing’s excel file output for every nascent spot over whole tissue. To make bar plots of exact number of nascent transcripts for each condition – wild type or RARE mutant, an average of 6-8 tail sections was plotted.

### Cell counts for each ROI

In the neural tube, the cells are densely packed together which makes it very hard to automate their counting. Conventional and Deep Learned algorithms have a hard time separating out cells and even manually it is difficult to go through each z-slice and count. To counter this problem. A sum projection file was generated using the DAPI channel for each tissue section. A small region where there is a lower density of cells was outlined, cells in that region were counted, and the integrated DAPI intensity per nuclei for this region was made. A ratio of integrated DAPI intensity/cell was used to roughly estimate the number of cells in the ROIs marked in each section. (Integrated DAPI Intensity of each ROI divided by the ratio of Integrated DAPI intensity/cell, to equal the number of cells present within the ROI). This was done for all tail section in all wild type and RARE mutants. For each tissue section, the exact number of nascent transcripts was divided by the corresponding cell counts for each ROI, and these values were used to plot the boxplots for the exact number of nascent transcripts per cell count.

### Statistical analysis of data

The nascent transcripts datasets were assessed for normality, and it was observed that the samples are bell-shaped. Hence, the non-parametric test Mann-Whitney (also called Wilcoxon rank-sum) test was performed, and all RARE mutant samples were compared against wild type for each probe set. Asterisks (*) over plots were used to denote significance. To calculate how range in nascent transcript expression was changing for each gene, variance was calculated by using a parametric F-test. Any significant change in variance (p-value < 0.05) compared against wild type samples was denoted by a star (*).

## Acknowledgements

The authors thank members of the Krumlauf laboratory and Viraj Doddihal, Vivekanandan Ramalingam, and Khyati Dalal for helpful discussions and feedback on the project and manuscript; Cindy Maddera for help in training the Deep Learning data sets and trouble-shooting imaging challenges; Youngwook Ahn, for generating the *DE* and *DE-B4U RARE* mutant mouse lines; and Heidi Monnin, Haley Tansey and Ian Castillo-Jolly collecting data on litter sizes and husbandry for mouse lines that are a challenge to maintain. We especially thank the Microscopy, Imaging and Big Data; Laboratory Animal Services; Histology; and Sequencing and Discovery Genomics technology facilities at Stowers Institute for their help in supporting this project. This work was conducted to fulfill, in part, the requirements for Z.A.’s degree as a PhD student with the Kansas University Medical Center, and we would like to thank the committee members for their insightful comments and feedback throughout.

## Competing interests

The authors declare no competing or financial interests.

## Author contributions

**Conceptualization** Z.A. and R.K.; **Methodology** Z.A., S.M., B.S., J.L., J.U., R.K.; **Software** Z.A., J.L., S.M., C.W., J.U., B.S.; **Validation** Z.A., J.L.; **Formal analysis** Z.A., J.L., S.M., J.U., B.S., A.P.; **Investigation** Z.A. J.L., B.S.; **Resources** C.N., Microscopy, Imaging and Big Data; Laboratory Animal Services; Histology; and Sequencing and Discovery Genomics technology facilities at Stowers Institute; **Data Curation** Z.A., B.S., J.L., J.U.; **Writing – original draft preparation** Z.A., R.K.; **Writing – review and editing** Z.A., R.K., J.L., B.S., C.N.; **Visualization** Z.A., J.L., J.U.; **Supervision** Z.A., B.S., J.U., B.K., R.K.; **Project administration** Z.A.; **Funding acquisition** This work was supported by funds from the Stowers Institute for Medical Research to R.K. (grant no: 1001) and to the Microscopy and Imaging technology center.

## Data availability

Original data underlying this manuscript can be accessed from the Stowers Original Data Repository at [http://odr.stowers.org/websimr/]

---

**Fig. S1.**
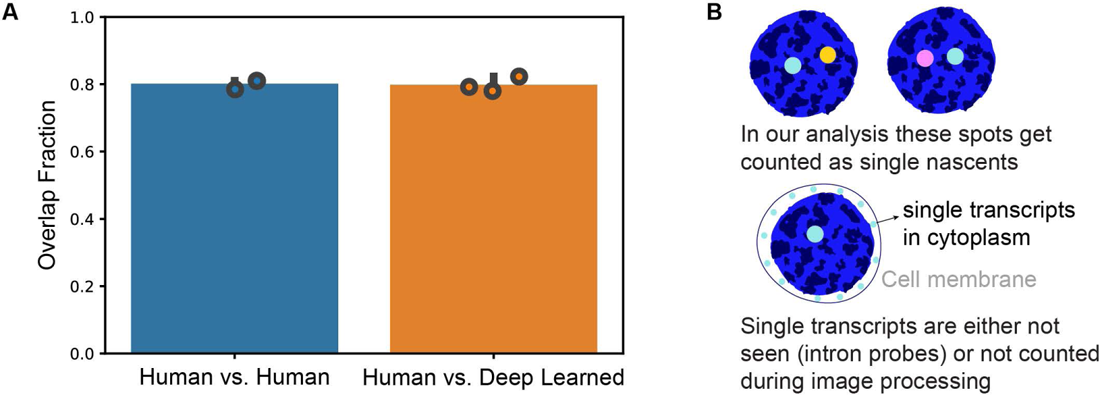
Reproducibility in calling spots of nascent transcripts. **A** Comparing accuracy of DL against manual markings of spots. **B** Cell schematics to indicate what doesn’t get counted as co-localized using Deep learning and a diagram to show that single transcripts in the cytoplasm do not get counted.

**Fig. S2.**
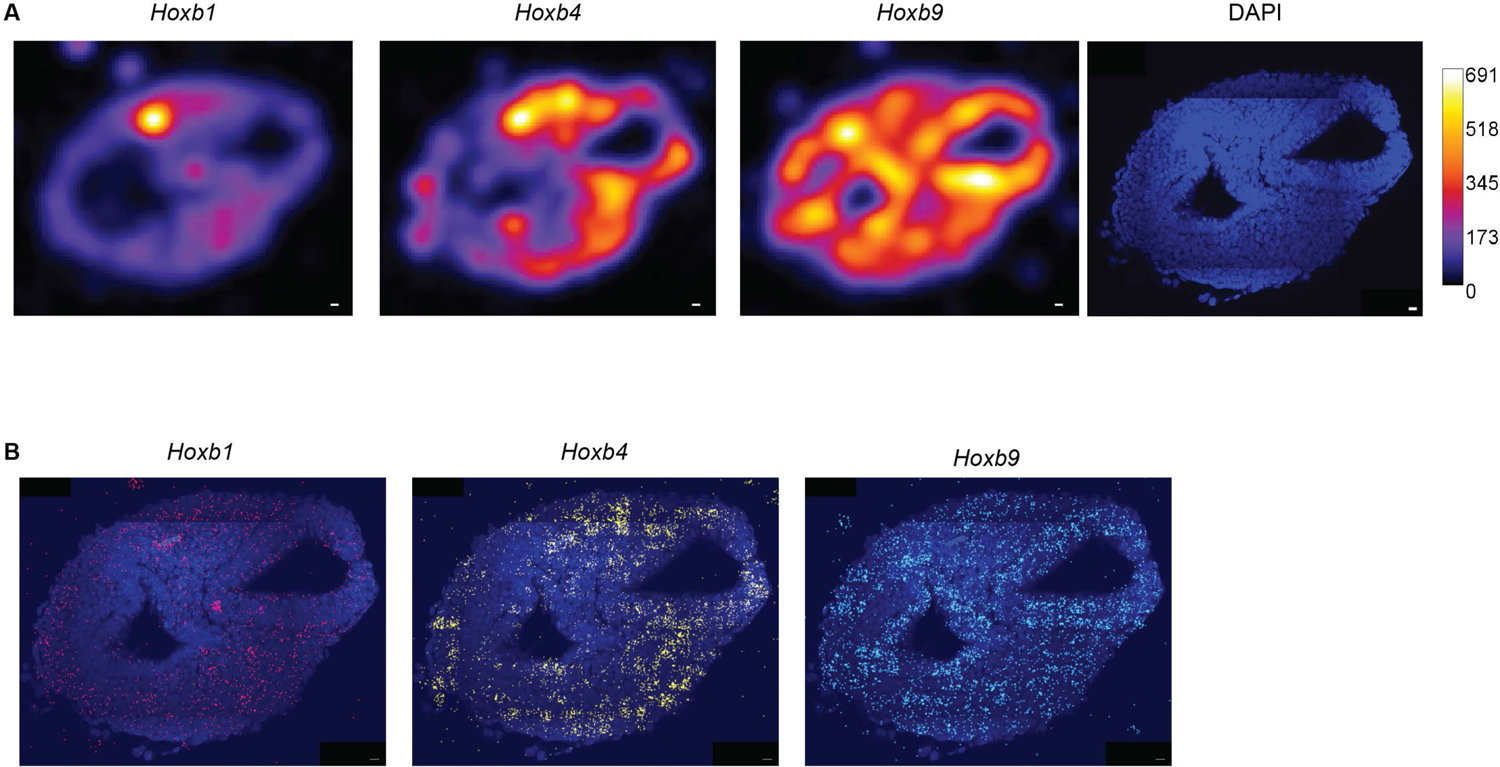
Intensity plots and spot fitting of nascent transcripts overlapping with DAPI staining. **A** Intensity plots to visualize regions of high or low nascent transcripts using fire LUT; white/yellow indicate higher numbers of nascent transcripts while blue/purple indicates lower numbers of nascent transcripts. **B** Spot fitting algorithm to visualize nascent transcripts overlapping with DAPI stained nuclei in the mouse tail.

**Fig. S3.**
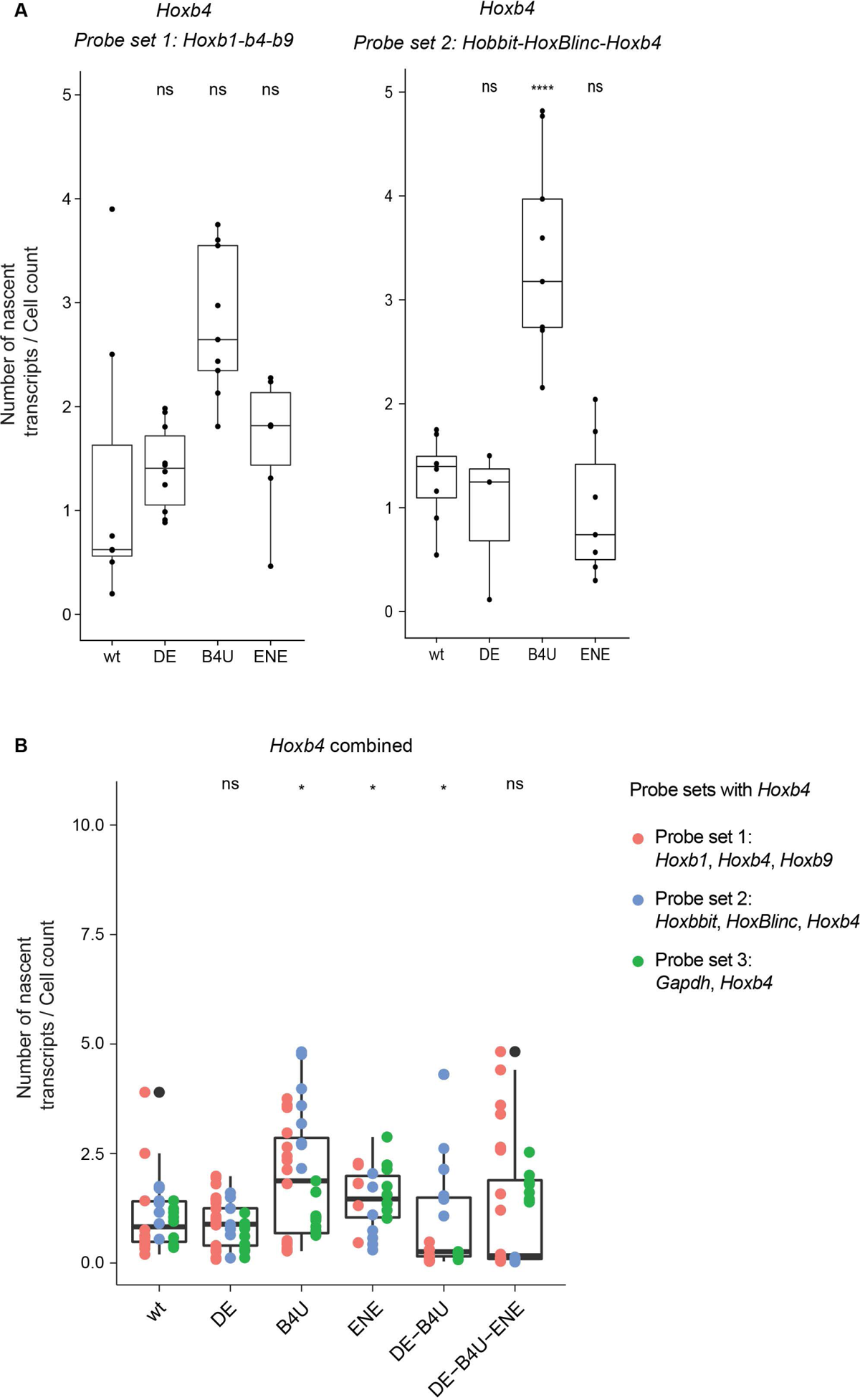
Quantification of total number of Hoxb4 nascent transcripts/cell in wild type and mutant embryos calculated as an average of data from multiple near adjacent tail sections. **A** Box plot of average levels of nascent Hoxb4 expression measured using with two different probe sets on a series alternate tissue sections from single embryo of wild type and mutant mouse lines. **B** Box plot of average levels of nascent Hoxb4 expression calculated by combining data from three different probe sets and multiple embryos. Each color represents a different probe set. Probes for the genes in each set are indicated at the right.

**Fig. S4.**
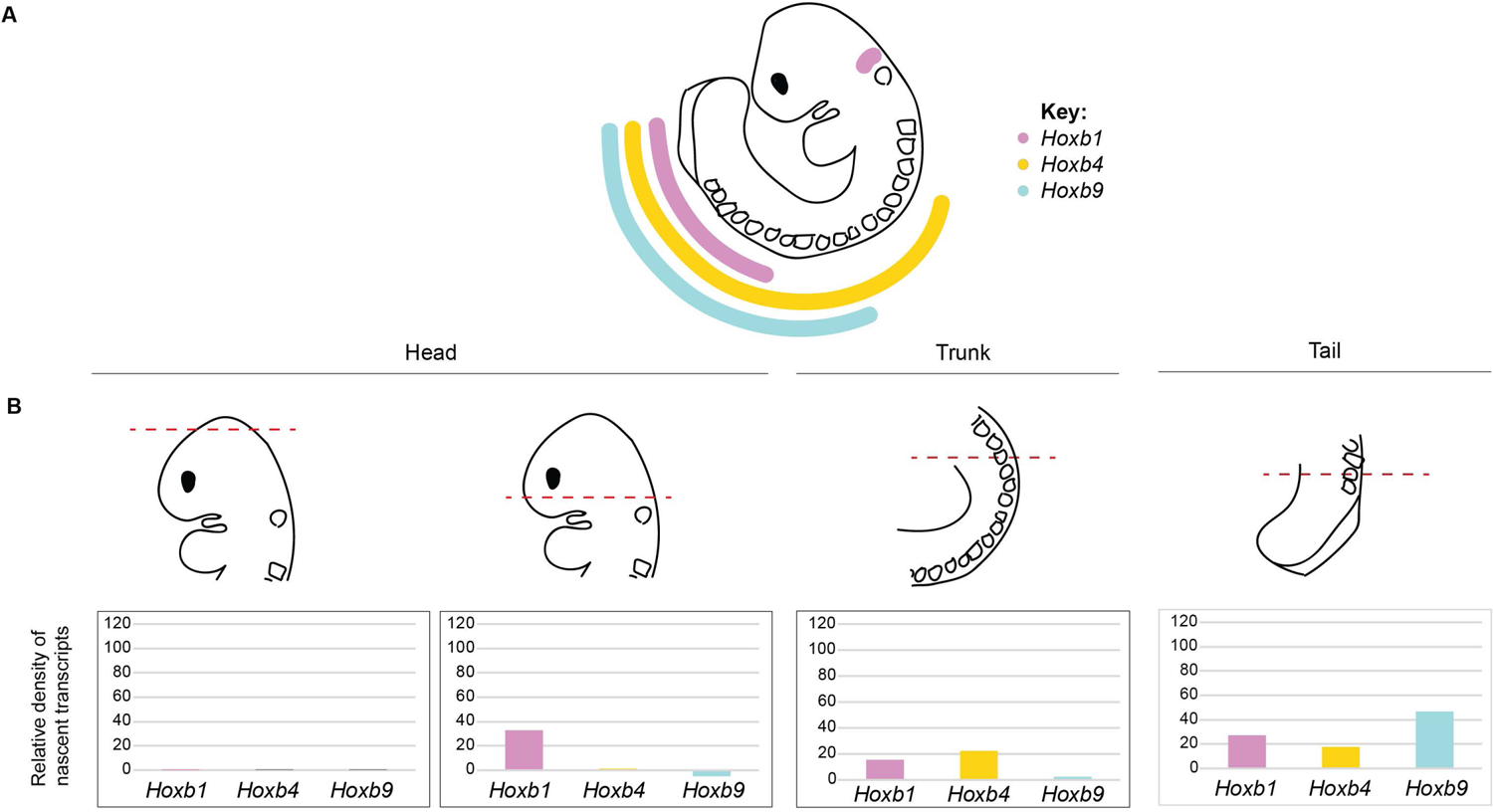
Comparison of detected Hoxb nascent transcripts against known expression. **A** Schematic illustrating known A-P expression patterns for steady state levels of *Hoxb1*, *Hoxb4*, and *Hoxb9* coding genesdetected by conventional colorimetric in situs in a 9.5 dpc mouse embryo. **B** At the top, a schematic of transverse sections (indicated by dotted line) at different A-P levels through neural tube of a 9.5 dpc embryo. Below each section are representative bar graphs indicating the relative densities in patterns of nascent transcription for *Hoxb1*, *Hoxb4* and *Hoxb9*. The shifts in level along the A-P axis mirror those illustrated in panel A, with *Hoxb4* and *Hoxb9* missing in anterior regions and elevated in posterior regions.

**Fig. S5.**
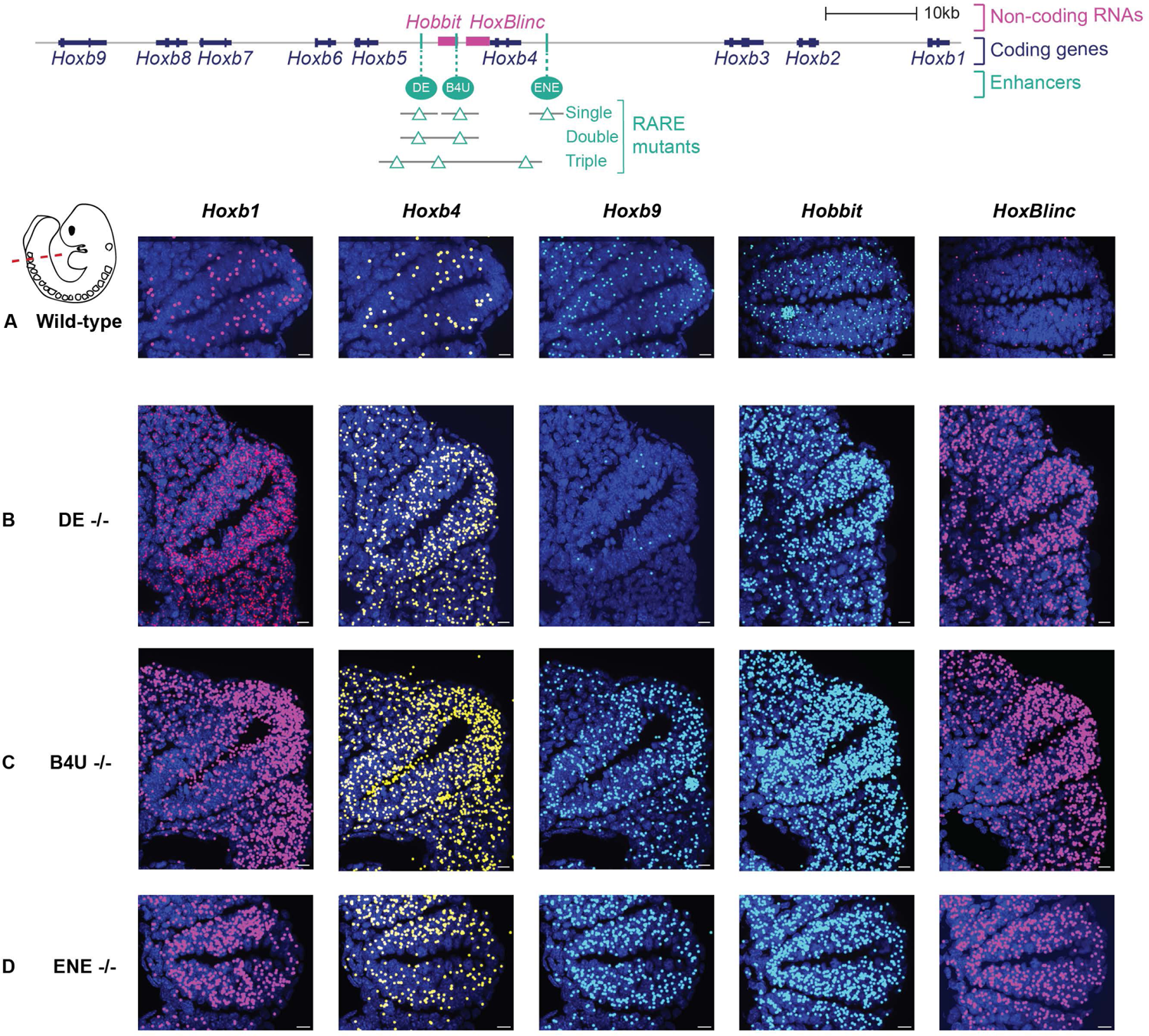
Visualizing changes in levels of nascent transcription of Hoxb coding and non-coding genes in single mutants of three different shared RARE enhancers. **A** Diagram depicting a series of single and compound mutants generated in RAREs of three shared enhancers present in the center of the *Hoxb* cluster. **B** Images of nascent transcripts for *Hoxb* coding and non-coding genes in wild type and the series of single RARE mutant embryos. The nascent transcripts detected by DL were spot fitted to increase the size of the spots for better visualization over the neural tube.

**Fig. S6.**
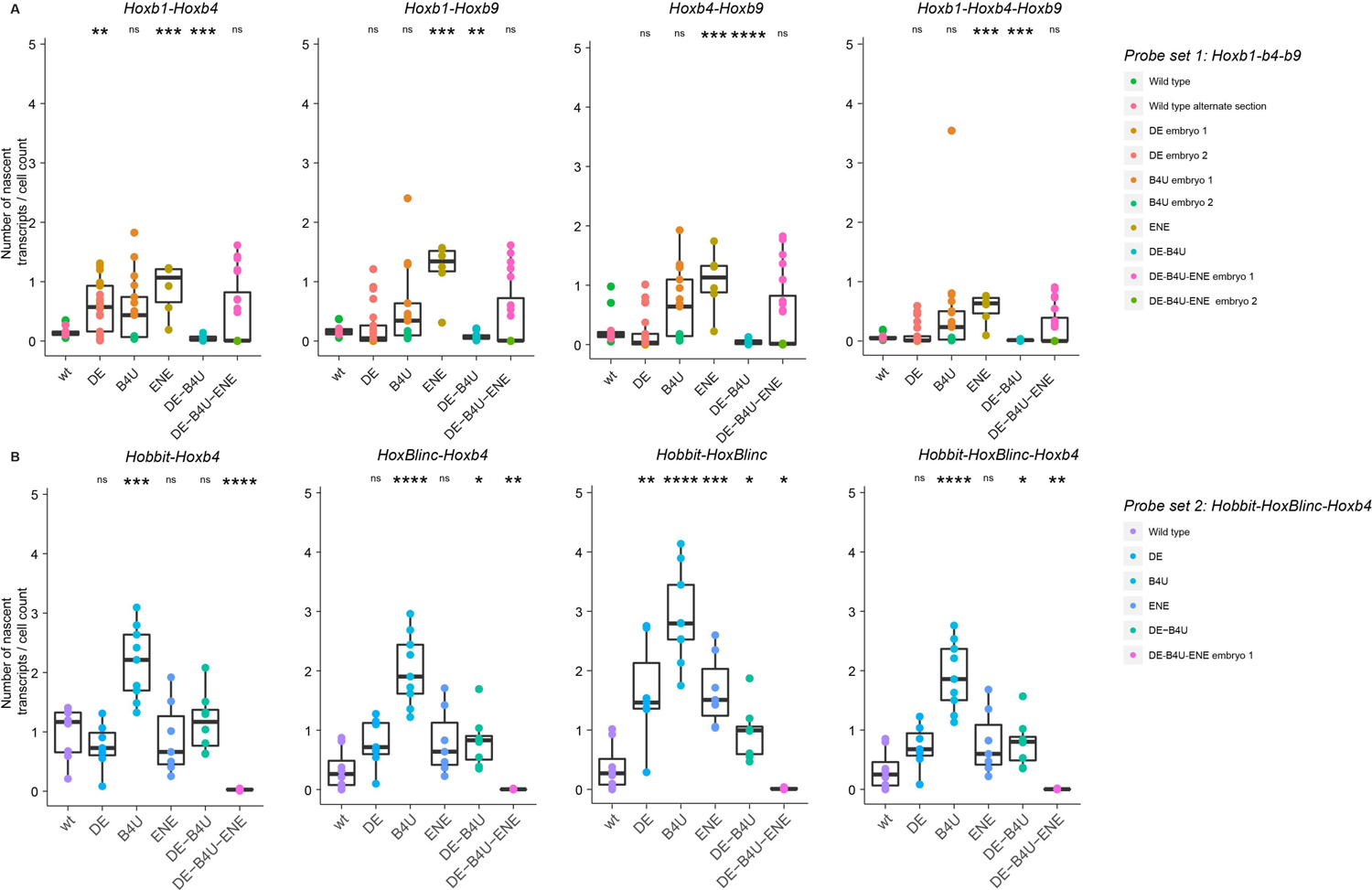
Quantifying patterns of co-localization of nascent transcription in wild type and mutant embryos. Box plots show the number of co-localized double and triple combinations of nascent transcripts/cell for *Hoxb* the coding (**A**) and non-coding (**B**) genes. Values are calculated as an average of combined data from multiple near adjacent tail sections (7-9) of multiple embryos. Each box in A and B represents the 75th (top line) and 25th (bottom line) percentile while the middle line represents the median expression.

**Fig. S7.**
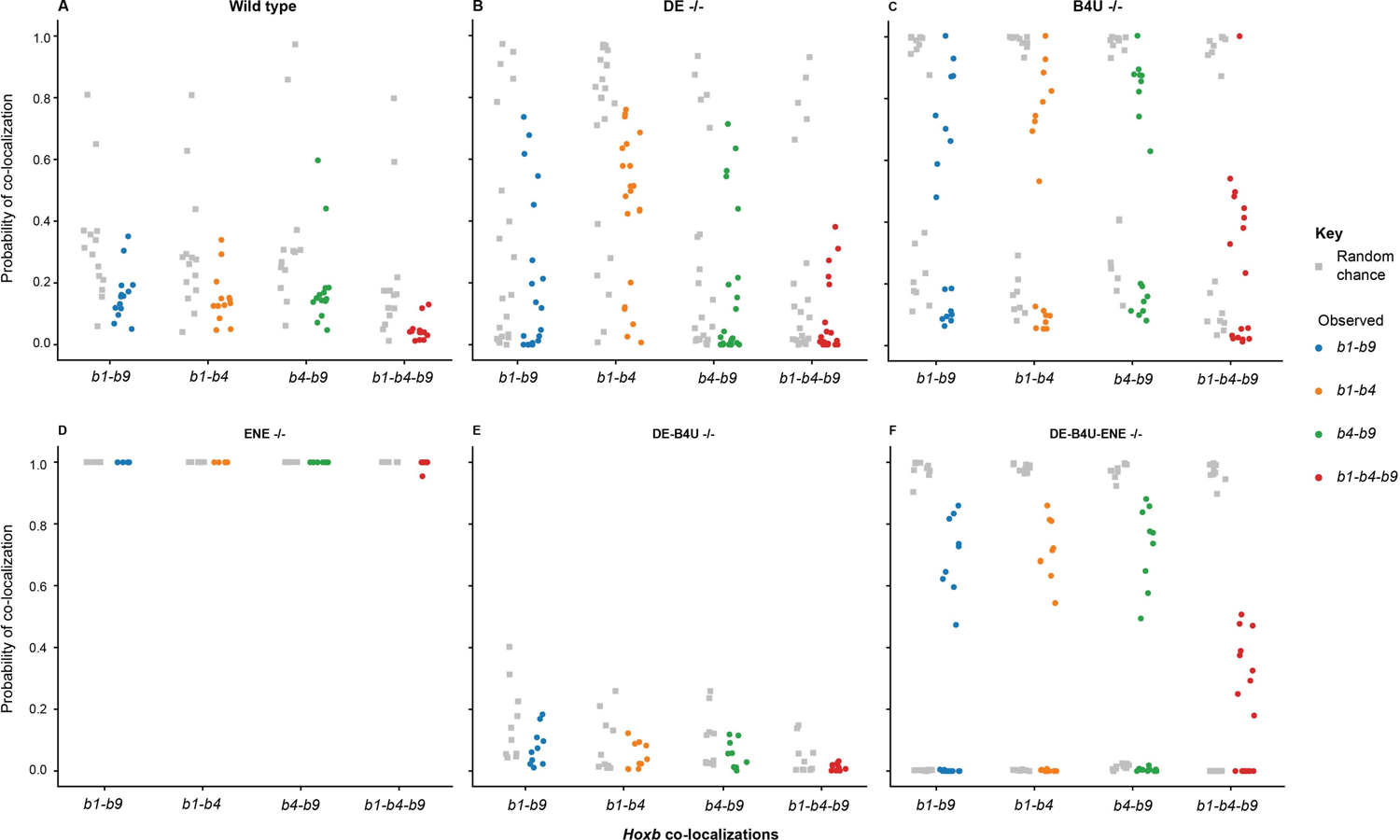
Observed and random probabilities for co-localization of Hoxb nascent transcripts. **A-F** Probabilities of co-localization by random chance (grey) next to observed co-localization probabilities (colored) for coding *Hoxb* genes in wild type and RARE mutants. Random chance of co-localization was calculated based on the numbers of single transcripts expressed within a defined area. The random values are compared against what was actually observed in single and compound RARE mutants.

**Fig. S8.**
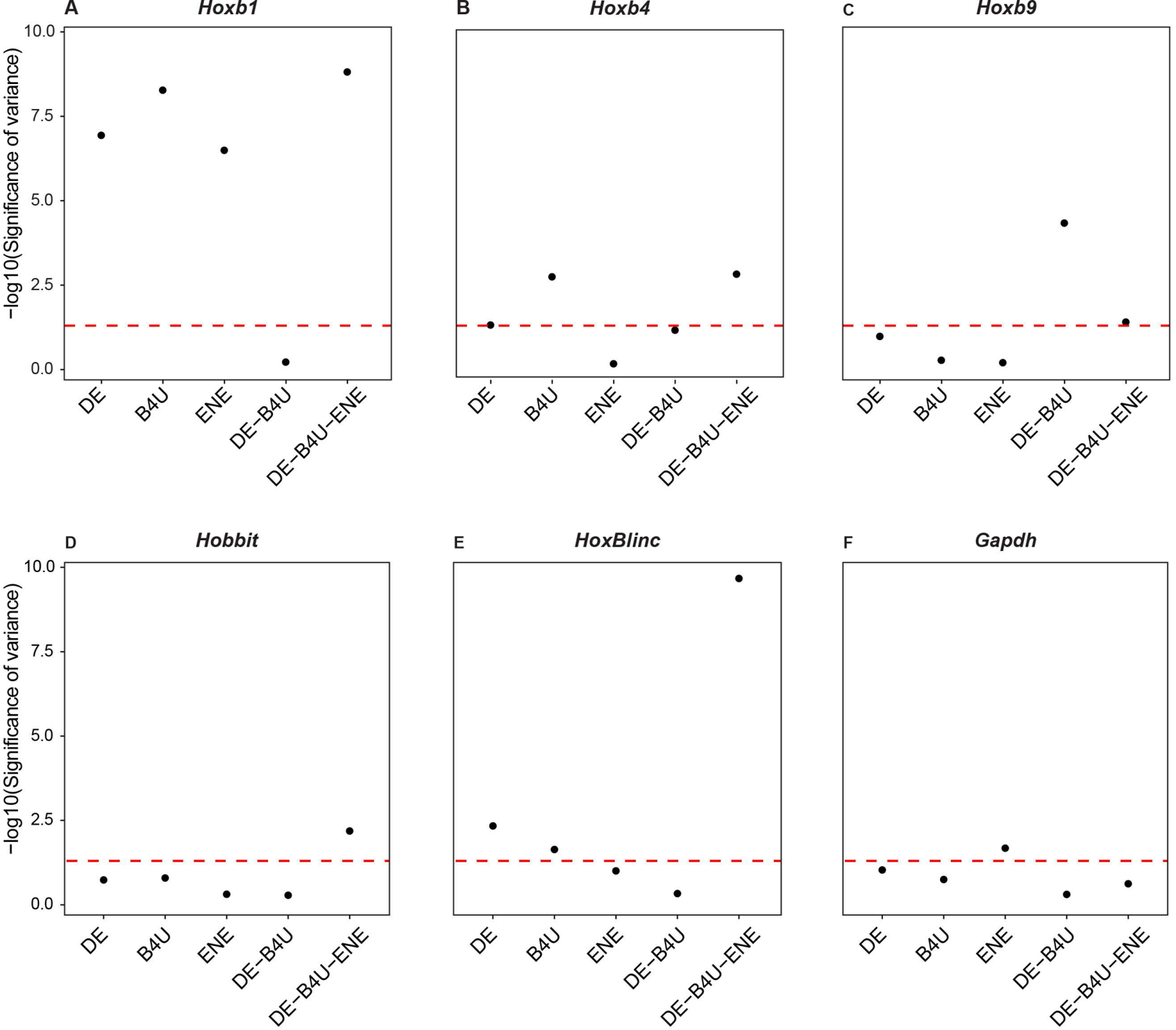
Variance plots for the difference in nascent transcripts in RARE mutants compared to wild type embryos. **A-F** Dot plots representing significance of variance in expression of *Hoxb* transcripts in RARE mutants compared to wild type. Red line represents p-value of 0.05 and thus any dot above the line is statistically significant for increased variance in expression of nascent transcripts among tissue sections compared to wild type.

**Table S1.** List of smFISH probes and their corresponding fluorophore. Three probe set combinations were used so genes could be imaged simultaneously without needing to spectrally unmix signals. Probe set 1 combined: *Hoxb9*-647, *Hoxb1*-570, and *Hoxb4*-488, Probe set 2: *Hobbit*-647, *HoxBlinc*-555, and *Hoxb4*-488, and Probe set 3: *Gapdh*-670 and *Hoxb4*-488.

| Genes | Labelled with fluorophore |
| --- | --- |
| *Hoxb1* exon | Quasar 570 |
| *Hoxb4* exon | Atto 488 |
| *Hoxb4* intron | AF 647 |
| *Hoxb9* intron | AF 647 |
| *Hobbit* | AF 647 |
| *HoxBlinc* | AF 555 |
| *Gapdh* | Quasar 670 |

**Table S2.**
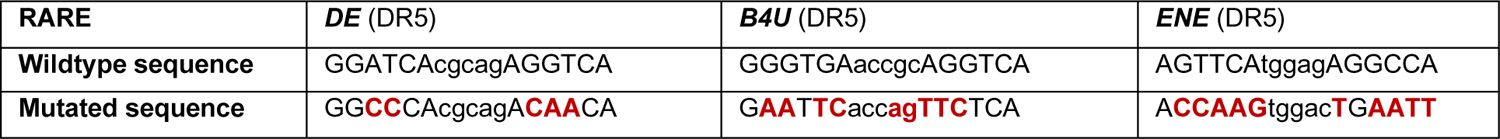
The specific mutations generated in the endogenous *DE*, *B4U* and *ENE RAREs* of the *Hoxb* cluster. The mutated sequences represent base pair substitutions (indicated in red) that disrupt RAR/RXR binding sites but maintain the same spatial distances between these and other cis-elements in the endogenous genomic locus. The sequences of the direct repeats (DR) in these RAREs are indicated in capital letters.

**Table S3.**
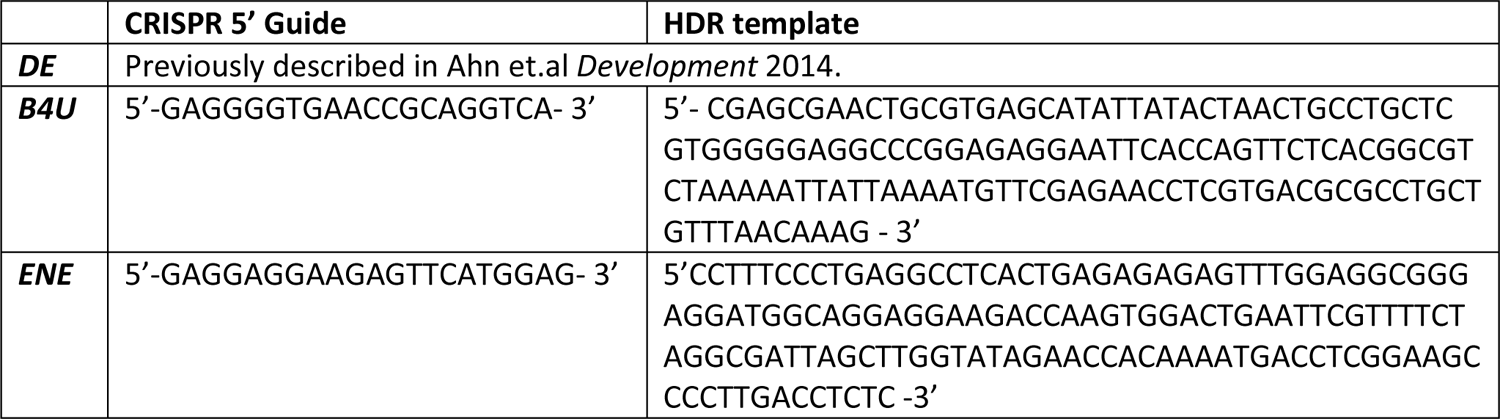
List of specific CRISPR guide and Homology Directed Repair (HDR) templates used to generate the *DE*, *B4U* and *ENE* RARE mutants in the endogenous *Hoxb* cluster. The sequences of CRISPR guides were ordered as oligos and the HDR templates were ordered as ultramers (IDT) and used to generate the RARE mutants in the Stowers F1 mouse strains. The *DE-B4U* double mutant was generated by microinjections of *B4U* guides in *DE-RARE* mutant animals, and the *DE-B4U-ENE* triple mutant was generated by microinjections of *ENE* guides in *DE-B4U RARE* mutant animals.

## Notes

### Competing Interest Statement

The authors have declared no competing interest.

